# Receptor dynamics and kinetic discrimination contribute to selectivity of β_1_-adrenoceptor antagonists

**DOI:** 10.64898/2026.11.10.765661

**Authors:** Qingchao Chen, Jillian G. Baker, Xin Xu, Tony Warne, Benjamin T. Porebski, Patricia C. Edwards, Gopal P. Jadhav, Christophe Fromont, Stuart J. Mundell, Philipp Holliger, Peter M. Fischer, Barrie Kellam, Shailesh N. Mistry, Stephen J. Hill, Robin A Corey, Christopher G. Tate

## Abstract

Human β_1_-adrenoceptor (hβ_1_AR)-selective antagonists are widely used to suppress cardiac adrenergic signalling in cardiovascular disease with limited but still clinically relevant inhibition of hβ_2_AR functions in airway and vascular smooth muscle. However, achieving strong subtype selectivity remains challenging because the orthosteric binding sites (OBS) of hβ_1_AR and hβ_2_AR are highly conserved. Here we studied the molecular mechanisms for three highly subtype selective hβ_1_AR antagonists, CGP20712A, NDD713 and NDD825. Three cryo-EM structures were therefore determined for each of the ligands bound to an inactive state of turkey tβ_1_AR in complex with an engineered nanobody fiducial (Nb60.5) to facilitate image processing. There were striking differences in how each antagonist bound to the receptor compared to what was observed in previous antagonist-bound tβ_1_AR and hβ_2_AR structures; these included non-canonical binding of the oxypropan-2-olamine (secondary amine and β-hydroxyl) cores of CGP20712A and NDD825 and the opening of a cleft between transmembrane helices H4 and H5 adjacent to the OBS. Mutagenesis of residues in the extended OBS that differ between hβ_1_AR and hβ_2_AR only partially explained the subtype selectivity. Subsequent kinetic analyses showed that all three ligands associated more rapidly with hβ_1_AR than hβ_2_AR, whereas dissociation effects were ligand dependent. Reciprocal receptor chimeras identified the hβ_1_AR extracellular vestibule as a key factor for fast ligand association. MD simulations further revealed that the cleft opening between H4 and H5 where CGP20712A and NDD825 insert is energetically feasible in hβ_1_AR, but is energetically inaccessible in hβ_2_AR. Together, these results establish that hβ_1_AR selectivity for CGP20712A and NDD825 arises from coordinated effects of OBS recognition, the ligand-entry pathway and receptor dynamics.

## Introduction

β-adrenoceptors (βARs) are class A G protein-coupled receptors (GPCRs) that mediate physiological responses to the catecholamines adrenaline and noradrenaline. The three βAR subtypes, β_1_AR, β_2_AR, and β_3_AR, can all activate G_s_-dependent cyclic AMP signalling, but differ in their tissue distribution and physiological functions^1^. β_1_AR mainly controls heart rate and contractility^2^, β_2_AR regulates smooth muscle relaxation in airways and blood vessels^3–5^ and β_3_AR activation predominantly stimulates lipolysis and thermogenesis in adipose tissue^6^. Subtype-selective ligands are therefore required to specifically modulate the receptors in disease states while minimising effects in other tissues.

Antagonists of hβ_1_AR, often called β-blockers, are commonly used to treat cardiovascular disorders such as heart failure, hypertension, ischemic heart disease, and arrhythmias^7^. Selective inhibition of cardiac hβ_1_AR is highly desirable because concomitant blockade of pulmonary hβ_2_AR can lead to bronchoconstriction and reduce the effectiveness of hβ_2_AR agonists, which is critical in patients who have both cardiovascular and respiratory diseases^8–10^. Although several β-blockers are characterised as cardioselective, their selectivity is limited^11^ and generally selectivity decreases as drug concentration increases^12^. Developing antagonists with enhanced hβ_1_AR selectivity could maintain cardiovascular effectiveness while reducing unwanted hβ_2_AR side-effects, but this is challenging because hβ_1_AR and hβ_2_AR are structurally very similar (Fig. 1a; in the transmembrane regions, 67% amino acid residue identity and RMSD 0.64 Å over 257 Cα atoms) and their OBSs are virtually identical (Fig. 1b). Previous crystal structures of tβ_1_AR^13–16^ showed that antagonists occupy the same position in the OBS and form interactions very similar to those in hβ_2_AR^17–19^. In particular, βAR agonists and antagonists have a shared polar core based on a secondary amine and a β-hydroxyl group that form conserved interactions with Asp^3.32^ and Asn^7.39^ (Fig. 1c; superscripts refer to the Ballesteros-Weinstein numbering convention^20^).

**Figure 1.**
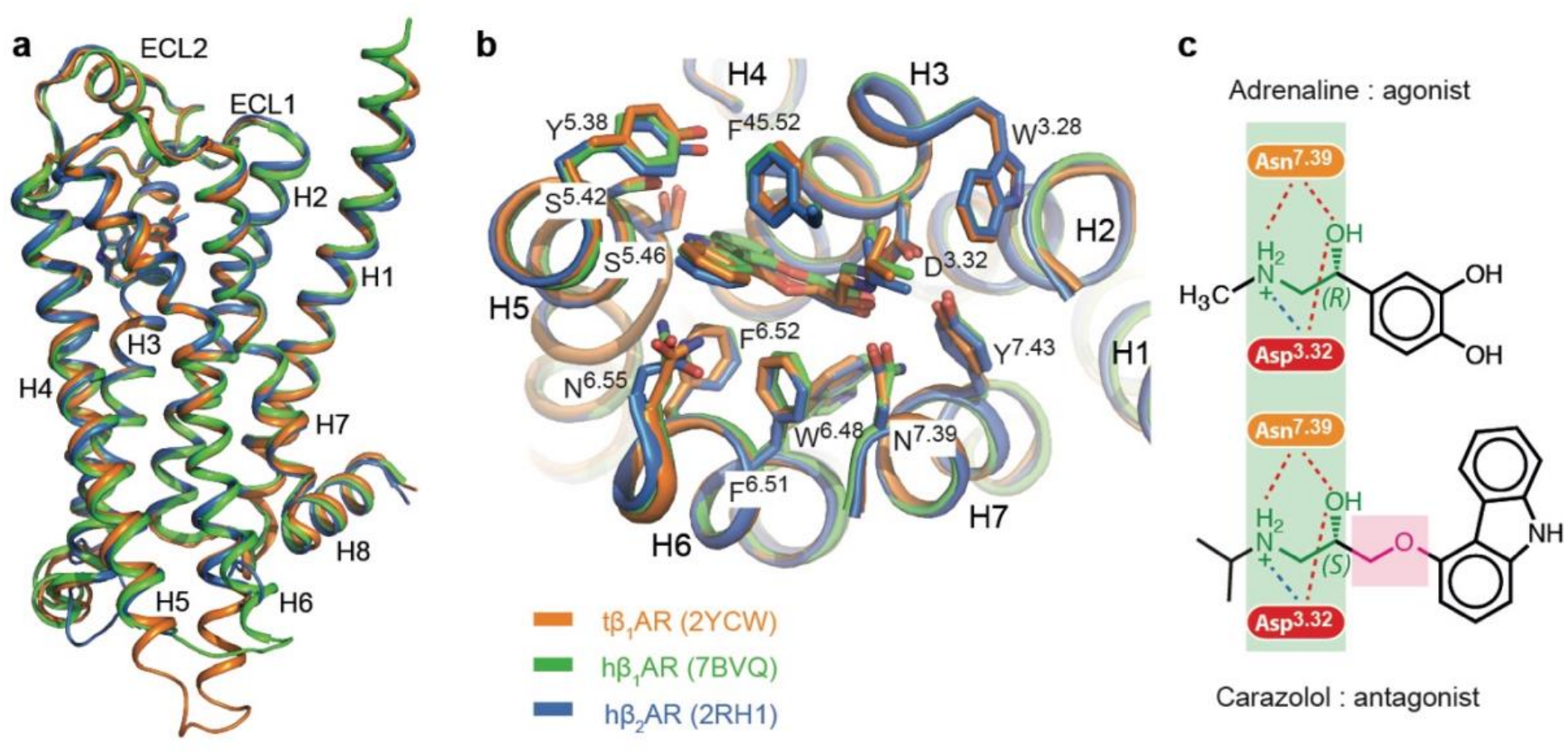
Similarities between the orthosteric binding site structure in tβ_1_AR, hβ_1_AR and hβ_2_AR. **a**, Structure superposition of carazolol-bound tβ_1_AR (PDB 2YCW), hβ_1_AR (7BVQ) and hβ_2_AR (PDB 2RH1). **b**, Detailed view of the OBS from the alignment shown in **a**, viewed from the extracellular surface with ECL1-3 removed for clarity. Receptor side chains ≤ 3.9 Å from carazolol are shown as sticks. **c**, Structures of the agonist adrenaline and antagonist carazolol, highlighting the conserved binding mode of the secondary amine and β-hydroxyl in both ligands (green) and the oxymethylene linkage inserted to make an antagonist (red). Residues are labelled with the Ballesteros-Weinstein nomenclature_20_ with dashed lines depicting potential hydrogen bonds (red) and a potential salt bridge (blue)_25_.

Ligand selectivity in hβ_1_AR can be influenced by non-conserved residues in H2 and H7^21–23^, but swapping individual pocket residues between hβ_1_AR and hβ_2_AR does not always transfer the complete pharmacological profile between them^23,24^. Increasing evidence suggests that hβ_1_AR selectivity can also be determined before a ligand reaches its final bound state. For example, the agonist noradrenaline exhibits an 8-fold higher affinity at hβ_1_AR compared to hβ_2_AR even though noradrenaline binds in a virtually identical manner to both receptors^25^. Computational studies^26^ suggested that ligand binding kinetics could explain ligand selectivity at the βARs due to variations in the shape and charge of the extracellular vestibule, and subsequent experiments swapping residues in the entrance to the OBS supports this^25^. Similarly, the selectivity of the antagonist bisoprolol for hβ_1_AR is partly due to its slower dissociation from the receptor^27^. Thus, ligand binding kinetics play a crucial role in ligand selectivity.

Among reported βAR antagonists, CGP20712A^28^, NDD713^29^ and NDD825^29^ display particularly pronounced hβ_1_AR selectivity. CGP20712A is an established hβ_1_AR-selective antagonist^28^, whilst NDD713 and NDD825 were developed more recently as highly selective hβ_1_AR neutral antagonists with minimal affinity for hβ_2_AR and favourable preclinical pharmacokinetic and safety properties^29^. Despite these antagonists having >500-fold^28,29^ selectivity for hβ_1_AR compared to hβ_2_AR, the molecular basis for this remains unknown. As defining selectivity determinants could inform the rational design of next-generation hβ_1_AR-selective antagonists, we determined structures of tβ_1_AR bound to CGP20712A, NDD713 and NDD825.

## Results

### Engineering a tβ_1_AR-binding nanobody and cryo-EM structure determination of antagonist-bound tβ_1_AR-nanobody complexes

To define the molecular determinants of CGP20712A, NDD713 and NDD825 selectivity at hβ_1_AR, we first needed to determine structures of them bound to tβ_1_AR. For structure determination a thermostabilised tβ_1_AR receptor was used rather than hβ_1_AR which is less stable^30^ and thus may have led to poorly resolved structures. Previous attempts to determine an X-ray structure of CGP20712A-bound tβ_1_AR were unsuccessful (Warne & Tate, unpublished data) as crystals did not grow despite using thermostabilised tβ_1_AR constructs that routinely yielded crystals and structures bound to different antagonists^13–16,31,32^. Single-particle electron cryomicroscopy (cryo-EM) was therefore the method of choice for determining the new structures. However, the small size of tβ_1_AR (32 kDa of structured protein), in combination with limited extracellular features extending beyond the detergent micelle, posed a significant challenge for particle alignment during image processing; initial attempts to determine structures of antagonist-bound tβ_1_AR were therefore unsuccessful (results not shown). To address this, we engineered a nanobody that would bind to the intracellular surface of inactive state tβ_1_AR and provide a fiducial feature outside the detergent micelle for particle alignment. This approach was previously successful for the cryo-EM structure determination of antagonist-bound hβ_2_AR in complex with nanobody Nb60, which bound to the intracellular surface of inactive state β_2_AR^33^.

There are 19 amino acid residues in hβ_2_AR that make contact to Nb60 and 17 of these are identical in tβ_1_AR, with conservative changes at the other two positions (Phe264 and Lys267 in hβ_2_AR are Met281 and Arg284 in tβ_1_AR, respectively). However, Nb60 showed no detectable binding to purified tβ_1_AR in the detergent lauryl maltoside neopentyl glycol (LMNG; results not shown). Superposition of the structures of Nb60-bound hβ_2_AR and tβ_1_AR suggested that this was probably due to an α-helix in intracellular loop 2 (ICL2) in tβ_1_AR that was absent in hβ_2_AR and would clash with the Nb60 complementarity-determining region 1 (CDR1; Fig. 2a). To overcome this, CDR1 was shortened by one or two residues (deletion of Asn32 or Leu31+Asn32) and then two libraries totalling approximately 6.6 × 10^6^ CDR1 variants were generated (Fig. 2b). A single round of Deep Screening^34^, a novel method for screening millions of antibodies, identified several hundred variants with detectable receptor binding profiles in the sub-500 nM range. Six nanobody variants with diverse CDR1 sequences and affinity profiles were selected for further characterisation (Fig. 2b). Biolayer interferometry (BLI) was then used to measure the binding kinetics of all six purified nanobodies (Fig. 2c). The six Nb60 variants showed different association and dissociation profiles, resulting in a range of binding affinities for tβ_1_AR (11 nM to 295 nM; Extended Data Fig. 1a-f). Nb60.5 was chosen for structural studies due to its favourable binding kinetics and its ability to form a stable complex with antagonist-bound tβ_1_AR in detergent (Extended Data Fig. 1g).

**Figure 2.**
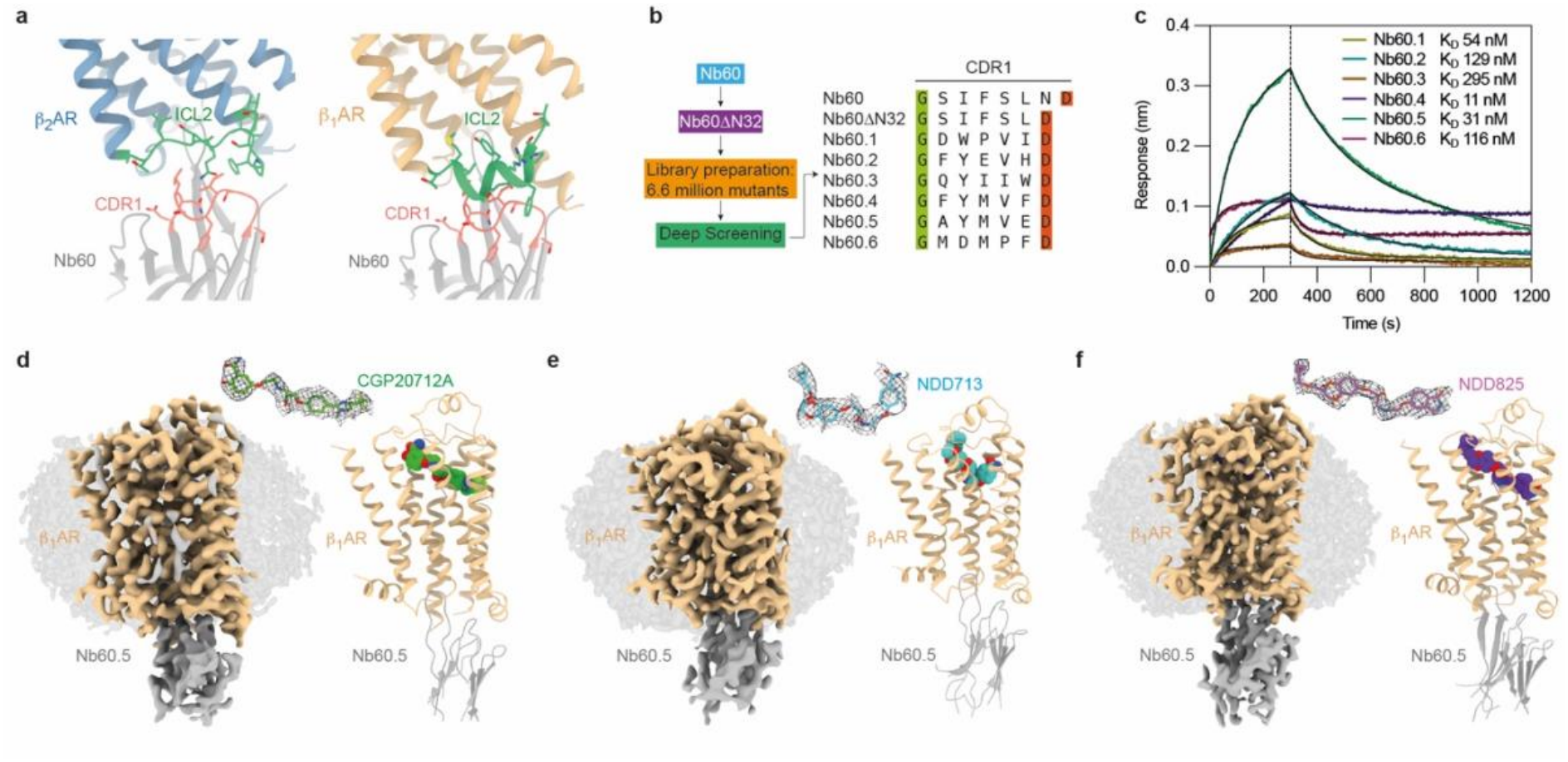
Cryo-EM structures of tβ_1_AR-Nb60.5 complexes bound to CGP20712A, NDD713 and NDD825. **a**, The Nb60-binding interface on hβ_2_AR (PDB 5JQH; left, blue) and the predicted clash between CDR1 (pink) and ICL2 (green) after alignment of the tβ_1_AR structure (PDB 2YCW; light brown, right) with the hβ_2_AR-Nb60 complex. Interface side chains are depicted as sticks. **b**, Workflow for nanobody engineering (left) and CDR1 sequences of the parental and selected nanobodies (right). The conserved flanking Gly and Asp residues are highlighted in green and orange, respectively. **c**, Biolayer interferometry (BLI) sensorgrams for binding of Nb60.1-Nb60.6 (100 nM concentration) to purified tβ_1_AR. Coloured traces show measured responses; black traces show global fits to a 1:1 binding model. The dashed line at 300 s marks dissociation onset. The K_D_ values were determined from BLI experiments performed at varying concentrations of nanobody (Extended Data Fig. 1a-f). **d– f**, Cryo-EM maps (left) and atomic models (right) of the tβ_1_AR-Nb60.5 complexes bound to CGP20712A (**d**), NDD713 (**e**) and NDD825 (**f**), at 3.0 Å, 3.3 Å and 2.8 Å resolution, respectively. Map densities are coloured as follows: light brown, tβ_1_AR; grey, Nb60.5; detergent micelle, pale grey. In the models, the ligands are shown as spheres (carbon atoms: green, CGP20712A; cyan, NDD713; purple, NDD825). The insets show each ligand as sticks within its cryo-EM density (mesh).

Structures were determined of an engineered tβ_1_AR that contained truncations to remove flexible regions in the N-terminus, ICL3 and C-terminus, mutations to improve folding (C116L, E130W), a thermostabilising mutation (M281F), removal of a palmitoylation site (C358A) and six mutations in ECL2 to change residues to those found in hβ_1_AR (D184A, D186S, P187D, Q188E, L190R, K191R). Cryo-EM of the antagonist-bound tβ_1_AR-Nb60.5 complex identified clearly the bound nanobody and therefore particle alignment was possible (Extended Data Fig. 2). Three cryo-EM structures were determined of tβ_1_AR bound to the antagonists CGP20712A, NDD713, and NDD825 (Fig. 2d-f) at global resolutions of 3.0 Å, 3.3 Å, and 2.8 Å, respectively (Extended Data Table 1, Extended Data Fig. 2a-c). The receptor transmembrane structures and ligand-binding pockets were well resolved and allowed us to define unambiguously the binding poses of all three ligands in the OBS (Extended Data Fig. 3a-c).

### Binding poses of hβ_1_AR-selective ligands

The structures of tβ_1_AR bound to CGP20712A, NDD713, and NDD825, were compared to tβ_1_AR bound to two archetypal ligands, the agonist isoprenaline^35^ (PDB: 2Y03) and the antagonist carvedilol (PDB: 4AMJ). The overall structures were all virtually identical to each other with RMSDs of 0.4-0.9 Å. All the ligands bound in the OBS (Fig. 3a). However, there were two striking features of how the hβ_1_AR subtype selective ligands bound that differed from what was expected for antagonists binding to tβ_1_AR, hβ_1_AR or β_2_AR. Firstly, the secondary amine and β-hydroxyl core of both NDD713 and NDD825 bound to tβ_1_AR in a non-canonical manner, even though the stereochemistry is identical to other antagonists *e.g.* carazolol (Fig. 1c) and carvedilol (Fig. 3d). NDD713 and NDD825 were developed through a traditional medicinal chemistry pharmacology screening programme without using molecular modelling for ligand design^36^. After development, modelling suggested they bound in a canonical manner with an extension protruding between H4 and H5^37^. The structures showed that both NDD713 and NDD825 do not bind in the manner that was originally anticipated, with the molecules rotated ∼180° (axis perpendicular to the membrane plane) in the OBS compared to docked ligand models reported in the original study^37^. It is also interesting that NDD713 binds in a kinked pose compared to NDD825, despite having a very similar structure (Fig. 3b,d). In all three structures, Asp121^3.32^ makes a potential hydrogen bond with the β-hydroxyl, but it makes a potential salt bridge to the secondary amine only with CGP20712A and NDD713 (Fig. 3d). None of the three antagonists make hydrogen bonds to Asn329^7.39^ as is found typically for other antagonists bound to tβ_1_AR, hβ_1_AR and hβ_2_AR (Fig. 1c, Fig. 3d). CGP20712A, NDD825 and NDD713 all extend the full length of the OBS, making contacts with H2 (Val102^2.65^ and/or Leu101^2.64^) at one end and with H5 (such as Ser21^15.42^, Fig. 3c) at the other end, as was also observed in the tβ_1_AR structures bound to bucindolol and carvedilol^15^. All the residues that make contact to either CGP20712A, NDD825 or NDD713 have been observed to make contacts before with other ligands in tβ_1_AR, hβ_1_AR and β_2_AR inactive state structures (Extended Data Fig. 4)^38^. However, there are some unusual omissions of contact residues. For example, none of the three antagonists studied here make contacts with Val125^3.36^ and Trp303^6.48^, which are observed in all other antagonist-bound βAR structures. Similarly, CGP20712A and NDD825 do not make interactions with Phe201^45.52^ and Asn310^6.55^, CGP20712A does not make contacts to Phe307^6.52^ and NDD713 does not make contacts to Ser215^5.46^. These comparisons highlight the unusual binding mode of these three antagonists.

**Figure 3.**
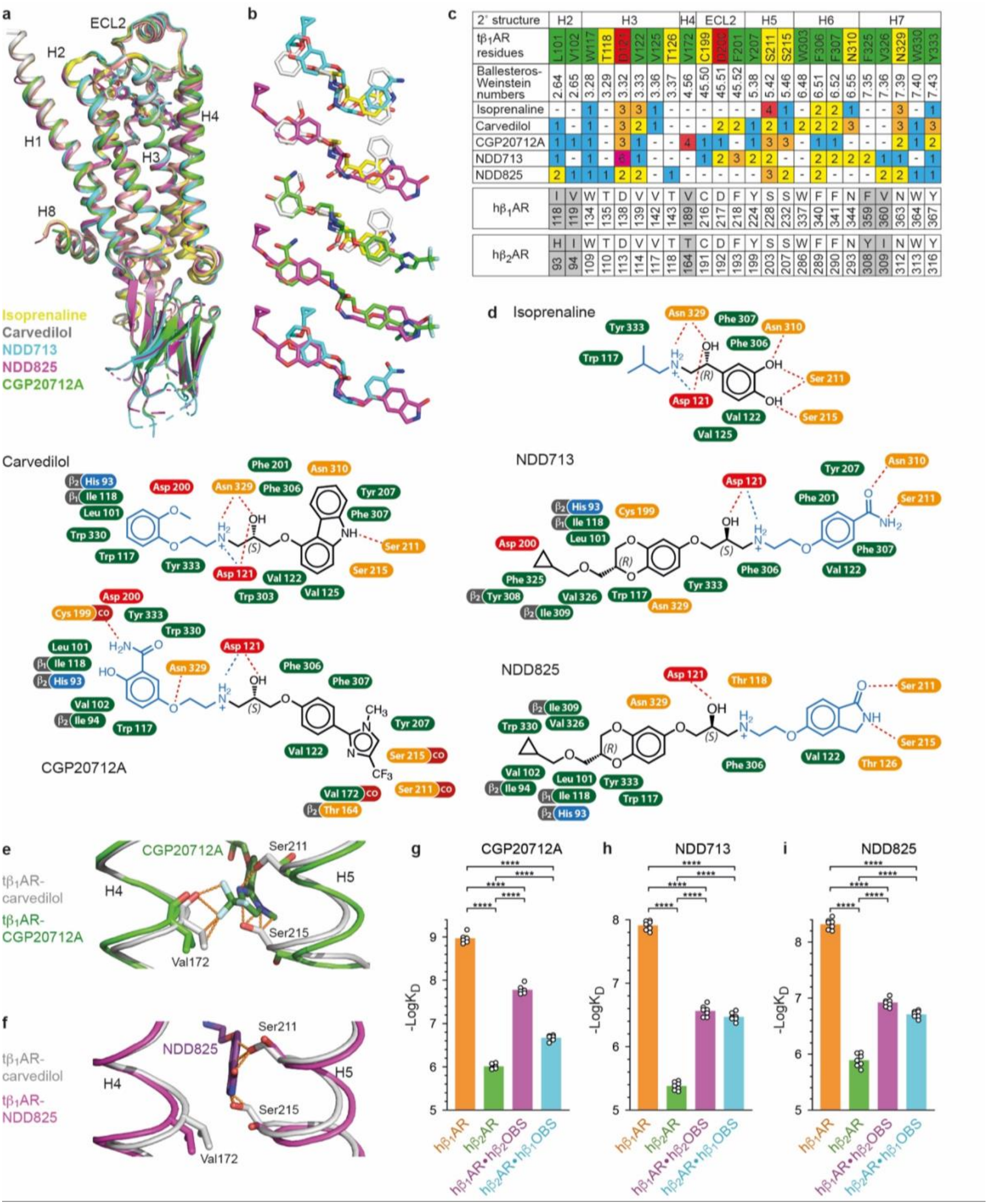
Binding modes of CGP20712A, NDD713 and NDD825 in tβ_1_AR compared with isoprenaline and carvedilol. **a**, Superposition of tβ_1_AR structures bound to isoprenaline (yellow; PDB 2Y03), carvedilol (grey; PDB 4AMJ), NDD713 (cyan), NDD825 (magenta) and CGP20712A (green), aligned on receptor Cα atoms. **b**, Ligands from the superposition in **a**, are shown as sticks in combinations of 2-3 for clarity. **c**, Amino acid contacts made between each ligand and tβ_1_AR (distance cut-off ≤3.9 Å). Residues are coloured by side-chain type (green, hydrophobic; yellow, polar; red, acidic) and the number of atom contacts is given (–, no contact). Equivalent residues of hβ_1_AR and hβ_2_AR are also shown; grey shading mark positions where hβ_1_AR and hβ_2_AR differ. **d**, Diagrams showing the interactions of each ligand with tβ_1_AR. Red dashed lines indicate potential hydrogen bonds and the blue dashed lines indicate potential salt bridges^25^. Interacting residues are shown as labels coloured as above. Grey ‘β_1_’ and ‘β_2_’ tags give the residues found at that position in hβ_1_AR and hβ_2_AR, respectively, where it differs from tβ_1_AR. **e-f**, tβ_1_AR bound to either CGP20712A (green) or NDD825 (magenta) showing the opening between H4 and H5 compared to tβ_1_AR bound to carvedilol (grey). Dashed orange lines represent clashes (distances ≤ 2.5 Å) between the ligands and the superposed structure of tβ_1_AR bound to carvedilol. **g-i**, Affinities (K_D_ values) for CGP20712A, NDD713 and NDD825, respectively, binding to hβ_1_AR (orange), hβ_2_AR (green), hβ_1_AR•hβ_2_OBS (purple) and hβ_2_AR•hβ_1_OBS (blue). Affinities are the mean ± SEM of 6-7 separate experiments, each performed 2-3 times (Extended Data Table 2); ****, *P* values <10^-10^ (see Extended Data Table 2 for values)

The second unusual feature observed in the structures has not been seen before with any other βAR ligand. Both CGP20712A and NDD825 showed an unusual extended mode of binding (Fig. 3b), with the trifluoromethylimidazolyl group of CGP20712A and the isoindoline-1-one moiety of NDD825 inserting between H4 and H5 (Fig. 3e,f; Extended Data Fig. 5a-c) that have moved apart by 2.8 Å to 2.5 Å (measured between Cα atoms in Val172^4.56^ and Ser215^5.46^), respectively. This opening is likely a consequence of clashes between CGP20712A and Val172^4.56^, Ser215^5.46^ and Ser211^5.42^, with NDD825 making clashes only with Ser215^5.46^ and Ser211^5.42^ (clashes defined as ≤ 2.5 Å between ligands and residues in tβ_1_AR bound to carvedilol superposed with the respective cryo-EM structures). In contrast to NDD825, the corresponding phenylcarboxamide moiety of NDD713 does not insert between H4-H5 as it adopts a kinked structure (Fig. 3b). The interface between H4 and H5 in the middle of the lipid bilayer is particularly interesting in hβ_1_AR and hβ_2_AR, because there is a significant difference between the receptors at position 4.56 (Val in hβ_1_AR, Thr in hβ_2_AR), with the Thr^4.56^Leu polymorphism in hβ_2_AR resulting in increased basal activity and E_max_^39^. In addition, this is the position where a negative allosteric modulator of hβ_2_AR binds^40^. Given that this region is important during receptor activation suggests that it is likely dynamic. Furthermore, these structural differences between hβ_1_AR and hβ_2_AR suggest that this region might also contribute towards subtype selectivity.

### hβ_1_AR ligand selectivity is determined partially by residues in the OBS

To investigate whether subtype selectivity was determined by residues directly surrounding the ligand, a systematic comparison was made of receptor-ligand contacts within the OBS (distance ≤3.9 Å). Most residues making ligand contacts in the tβ_1_AR structures are conserved across tβ_1_AR, hβ_1_AR and hβ_2_AR (Fig. 3c-d). The single residue that differs in the OBS of tβ_1_AR and hβ_1_AR is the conservative change of Leu101^2.64^ to Ile118^2.64^, respectively, that makes contacts to all three antagonists (Fig. 3c-d). There are five differences between hβ_1_AR and hβ_2_AR, with the residues Ile118^2.64^, Val119^2.65^, Val189^4.56^, Phe359^7.35^, and Val360^7.36^ in hβ_1_AR being replaced by His93^2.64^, Ile94^2.65^, Thr164^4.56^, Tyr308^7.35^, and Ile309^7.36^ in hβ_2_AR, respectively (Fig. 3c-d). To determine their role in ligand selectivity, each of these five residues in the OBS of hβ_1_AR were mutated to the equivalent residues in hβ_2_AR, making an hβ_1_AR receptor mutant with the OBS residues of hβ_2_AR (mutant hβ_1_AR•hβ_2_OBS). An analogous mutant was made in hβ_2_AR containing the OBS residues of hβ_1_AR (mutant hβ_2_AR•hβ_1_OBS).

Radioligand binding assays using the antagonist [^3^H]CGP12177 recapitulated the strong hβ_1_AR selectivity for CGP20712A, NDD713 and NDD825 over hβ_2_AR (Fig. 3g-i, Extended Data Table 2) with similar selectivity values (913-fold, 339-fold and 269-fold, respectively) to those published previously^28,29,41^. In the hβ_1_AR•hβ_2_OBS mutant, the affinity for each ligand decreased (Fig. 3g-i, Extended Data Table 2) as observed by an increase in apparent K_D_ by 15-fold, 22-fold and 25-fold for CGP20712A, NDD713 and NDD825, respectively. The reciprocal substitutions in the mutant hβ_2_AR•hβ_1_OBS improved ligand affinity of β_2_AR, but the magnitude of this rescue was ligand-dependent; affinity increases of 5-fold, 12-fold and 7-fold were observed for CGP20712A, NDD713 and NDD825, respectively (Fig. 3g-i, Extended Data Table 2). Nevertheless, CGP20712A, NDD713 and NDD825 retained selectivity for hβ_1_AR over hβ_2_AR•hβ_1_OBS to the magnitude of 200-fold, 28-fold and 41-fold respectively.

To define the role of individual hβ_1_AR residues in determining selectivity of the CGP20712A, NDD713 and NDD825, assays were performed on single mutants for each of the residues that differed between hβ_1_AR and hβ_2_AR in the OBS (Extended Data Fig. 6 and Extended Data Table 3). Binding of CGP20712A and NDD713 to hβ_2_AR was enhanced by the mutation H93I^2.64^ and the converse mutation in hβ_1_AR (I118H^2.64^) reduced binding; no further effect was seen after adding the additional four mutations in the hβ_2_AR•hβ_1_OBS mutant. Binding of NDD825 to hβ_2_AR was enhanced by both H93I^2.64^ and I94V^2.65^, and the converse mutations in hβ_1_AR (I118H^2.64^ and V119I^2.65^) reduced binding; no further effect was seen after adding the additional three mutations in the hβ_2_AR•hβ_1_OBS mutant. The results differ slightly when considering the effect of residues from hβ_2_AR introduced into the OBS of hβ_1_AR. The binding of both CGP20712A and NDD713 are reduced by the mutation I118H^2.64^, but also by an additional mutation F359Y^7.35^ and there was a significant additive effect of combining all five mutations in the mutant hβ_1_AR•hβ_2_OBS. Similarly, NDD825 binding to hβ_1_OBS was reduced by each of the mutations I118H^2.64^, V119I^2.65^ and F359Y^7.35^ and combining all five mutations in the hβ_1_AR•hβ_2_OBS mutant was additive.

These mutational data showed that the five non-conserved OBS residues play a significant role in subtype-selective affinity, but neither hβ_1_AR•hβ_2_OBS or hβ_2_AR•hβ_1_OBS recapitulated the corresponding wild-type receptor affinity. This suggested that features of the receptor beyond the OBS are necessary to account for complete hβ_1_AR selectivity of the ligands.

### The extracellular vestibule contributes to hβ_1_AR selectivity

The extracellular regions of hβ_1_AR and hβ_2_AR play an important role in the subtype selectivity of noradrenaline (norepinephrine) by affecting its rate of association^25^. To explore this possibility with the hβ_1_AR selective antagonists CGP20712A, NDD713 and NDD825, two reciprocal chimeras were generated in which the extracellular regions of hβ_1_AR and hβ_2_AR were exchanged, while the corresponding transmembrane cores, intracellular regions and the OBS residues were retained^25^. The mutant receptor hβ_1_AR_out_/hβ_2_AR_in_ had the β_2_AR transmembrane core and the hβ_1_AR extracellular region (Extended Data Fig. 7a); conversely, hβ_2_AR_out_/hβ_1_AR_in_ had the opposite combination (Extended Data Fig. 7b). Four stable cell lines were generated for competition-association experiments^42^, including wild-type hβ_1_AR, wild-type hβ_2_AR, hβ_1_AR_out_/hβ_2_AR_in_, and hβ_2_AR_out_/hβ_1_AR_in_. We performed competition-association experiments and analysed the resulting data using a kinetic model of competitive binding to determine ligand association (k_on_) and dissociation rates (k_off_), from which kinetic K_D_ values were calculated (Extended Data Figs. 8 and 9).

All three ligands showed significant kinetic selectivity for hβ_1_AR over hβ_2_AR. CGP20712A associated with hβ_1_AR 67 times faster and dissociated 26 times more slowly than with hβ_2_AR, resulting in a 1,747-fold difference in kinetic affinity (Fig. 4a-c). NDD713 showed an 818-fold kinetic affinity difference, mainly due to a 337-fold faster association rate, along with a modest 2.4-fold decrease in dissociation rate (Fig. 4d-f). In contrast, the 978-fold kinetic affinity difference of NDD825 came from both a 22-fold faster association rate and a 44-fold slower dissociation rate (Fig. 4g-i). Thus, the binding affinities of the three ligands are determined by both a faster on-rate and a slower off-rate, but these vary among them.

**Figure 4.**
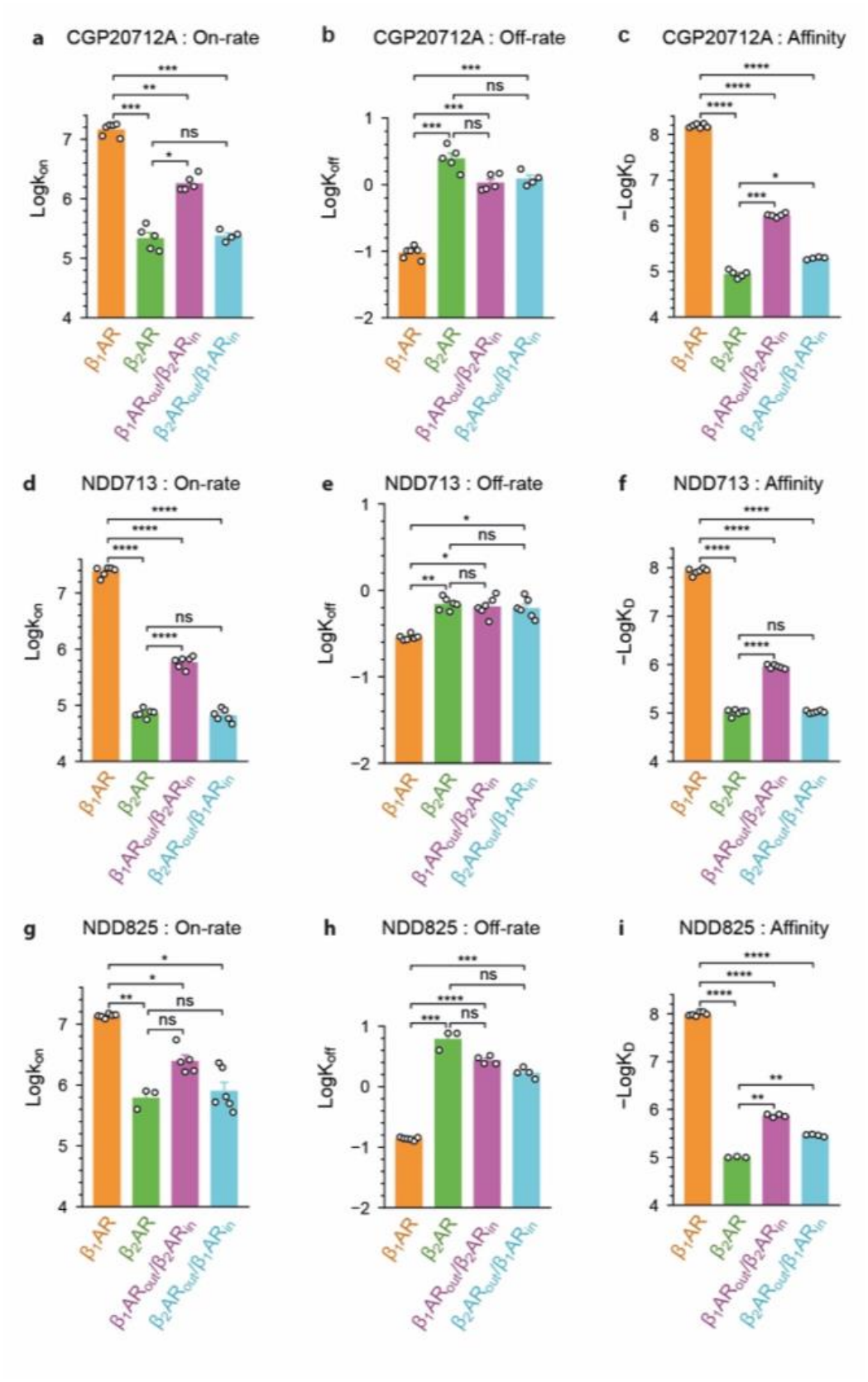
Association and dissociation rates for CGP20712A, NDD713 and NDD825. **a**-**c**, Association rate constants (log *k*_on_, a), dissociation rate constants (log *k*_off_, b) and affinities (−log *K*_D_, c) of CGP20712A at wild-type hβ_1_AR and hβ_2_AR and the chimeric receptors hβ_1_AR_out_/hβ_2_AR_in_ and hβ_2_AR_out_/hβ_1_AR_in_, determined by radioligand binding kinetics using [³H]dihydroalprenolol ([³H]DHA). **d**-**f**, The same parameters for NDD713. **g**-**i**, The same parameters for NDD825. Data are mean ± SEM of n = 3-6 performed in duplicate. Comparisons were tested using an unpaired two-tailed Student’s t-test (*\*\*\*\**, *P* ≤ 10^-9^; ***, *P* ≤ 10^-7^; **, *P* ≤ 10^-5^; *, *P* ≤ 10^-3^; ns, P > 10^-3^). All the data values and significance values are listed in Extended Data Table 4. Underlying data for ligand binding, association and dissociation are shown in Extended Data Figs. 8 and 9.

Changing the extracellular vestibule significantly affected ligand association. The hβ_1_AR_out_/hβ_2_AR_in_ increased the rate of association compared to hβ_2_AR by 8-fold for CGP20712A, 8-fold for NDD713, and 4-fold for NDD825 compared to β_2_AR (Fig. 4a,d,g). The reverse substitution had an even greater effect: hβ_2_AR_out_/hβ_1_AR_in_ slowed association compared to that for hβ_1_AR by 61-fold for CGP20712A, 367-fold for NDD713, and 17-fold for NDD825 (Fig. 4a,d,g). In fact, the association of all three ligands with hβ_2_AR_out_/hβ_1_AR_in_ was nearly the same as that with hβ_2_AR, despite retaining the hβ_1_AR transmembrane core. The swapping of extracellular regions also influenced ligand dissociation, particularly for CGP20712A and NDD825. In the mutant hβ_2_AR_out_/hβ_1_AR_in_ the rate of dissociation compared to hβ_1_AR was increased for CGP20712A and NDD825 by 13-fold and 12-fold, respectively, but only 2.2-fold for NDD713 (Fig. 4b,e,h). Conversely, in hβ_1_AR_out_/hβ_2_AR_in_ the rate of dissociation of CGP20712A and NDD825 was slowed by 2.3-fold and 2.2-fold, respectively, while having little effect on NDD713 (Fig. 4b,e,h). The extracellular region therefore plays a role in both ligand association and dissociation for CGP20712A and NDD825, while its effect on NDD713 selectivity is dominated by association.

In general, the kinetic effects of the extracellular vestibule exchanges led to significant changes in affinity. Compared to hβ_2_AR, hβ_1_AR_out_/hβ_2_AR_in_ showed 19-fold, 8.7-fold, and 7.4-fold higher kinetic affinity for CGP20712A, NDD713 and NDD825, respectively (Fig. 4c,f,i). In contrast, hβ_2_AR_out_/hβ_1_AR_in_ reduced affinities by 789-fold, 801-fold, and 342-fold, respectively (Fig. 4c,f,i). However, neither reciprocal chimera fully recapitulates the binding affinities for the receptor that donated its extracellular vestibule. Specifically, hβ_1_AR_out_/hβ_2_AR_in_ had an affinity that remained 90-fold to 133-fold lower than hβ_1_AR. Thus, the extracellular regions of hβ_1_AR contribute to the selectivity of CGP20712A, NDD713 and NDD825 but they are not sufficient alone to explain the high hβ_1_AR selectivity of the three antagonists.

### A transmembrane H4-H5 open conformation in hβ_1_AR

Amino acid residue differences between hβ_1_AR and hβ_2_AR in the OBS and extracellular region each accounted for part of the ligand selectivity for hβ_1_AR of CGP20712A, NDD713 and NDD825. However, the cryo-EM structures identified a potential dynamic component of ligand recognition at the tβ_1_AR H4-H5 interface. In the CGP20712A and NDD825 structures, a hydrophobic moiety extended into a cleft between H4 and H5 (Fig. 3e,f). The trifluoromethyl group of CGP20712A and the isoindoline-1-one group of NDD825 penetrated this interhelical cleft, creating a binding pose not observed in any other tβ_1_AR, hβ_1_AR and hβ_2_AR structures. We hypothesise that preferential sampling of an open interhelical conformation by hβ_1_AR could increase the probability of forming a productive binding intermediate capable of accommodating the extended ligand groups. Conversely, limited access to this conformation in hβ_2_AR might create an energetic barrier to ligand entry or insertion and hence lead to selectivity of ligand binding. To test this hypothesis, molecular dynamics simulations were used to probe the dynamics of the H4-H5 interface.

MD simulations with umbrella sampling (Extended Data Table 5) were used to calculated potentials of mean force (PMFs) for the separation of H4 and H5 in ligand-free (apo) hβ_1_AR and hβ_2_AR, using the centre-of-mass distance, q, between selected residues on the two helices (Fig. 5a). A total of 16 umbrella windows were centred from q = 0.8 to 1.12 nm, spaced at 0.021 nm intervals along the reaction coordinate. Initial conformations were extracted from unbiased apo simulations, meaning the input landscapes were sampled without applying any external biases. The reconstructed PMF extended to approximately q = 0.70 to 1.20 nm considering the sampled fluctuations in each window. We compared the computed free-energy profiles for H4-H5 separation in apo hβ_1_AR and hβ_2_AR with reference distances from inactive state structures of hβ_1_AR (PDB 7BVQ, q = 0.82 nm) and hβ_2_AR (PDB 2RH1, q = 0.86 nm), and the tβ_1_AR CGP20712A-bound structure (q* = 1.06 nm). The hβ_1_AR PMF contained a global minimum centred around q = 0.86 nm and a second, shallower, local minimum at q = 1.03 nm, close to q* (Fig. 5b). The H4-H5 separation observed in CGP20712A-bound tβ_1_AR (1.06 nm) therefore lies within a locally favourable region of the apo hβ_1_AR free-energy profile. By contrast, the hβ_2_AR PMF had a broad global minimum from q = 0.85-0.94 nm and rose monotonically towards q*, without a corresponding local minimum near the CGP20712A-bound separation (Fig. 5b). The bootstrap mean PMF at q* in relation to the global minimum for each system was 0.55 kcal/mol for hβ_1_AR and 1.34 kcal/mol for hβ_2_AR, suggesting a ΔΔG of 0.79 kcal/mol. These profiles therefore support the compatibility of the inserted CGP20712A pose with the inherent conformational dynamics of hβ_1_AR but not hβ_2_AR.

**Figure 5.**
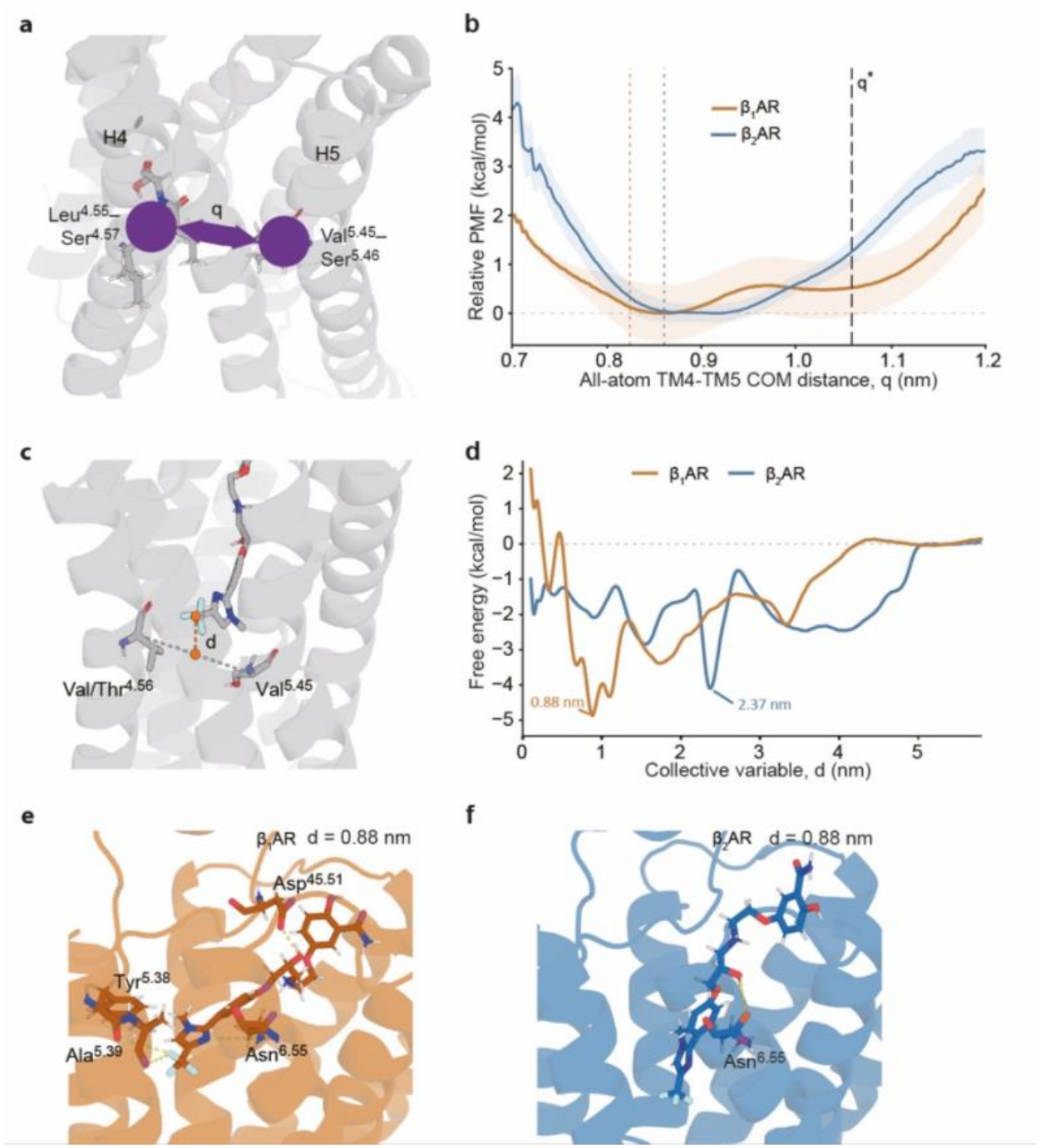
Molecular dynamics simulations of CGP20712A in β_1_AR and β_2_AR. **a,** Definition of the reaction coordinate q used for the umbrella-sampling simulations: the centre-of-mass distance between Leu^4.55^–Ser^4.57^ (residues 188-190 in hβ_1_AR and 163-165 in hβ_2_AR) on H4 and Val^5.45^–Ser^5.46^ (residues 231-232 in hβ_1_AR and 206-207 in hβ_2_AR) on H5 (purple spheres, arrow). The receptor is shown as a grey cartoon. **b,** Potentials of mean force for H4-H5 separation in apo hβ_1_AR (pale brown) and apo hβ_2_AR (blue), from 16 umbrella windows spaced at 0.021 nm along q, with windows seeded from unbiased apo simulations. Shaded areas are bootstrap 95% confidence intervals. Dotted vertical lines mark q in the inactive-state structures of hβ_1_AR (0.82 nm; PDB 7BVQ) and β_2_AR (0.86 nm; PDB 2RH1); the dashed line marks q* = 1.06 nm, the separation in the CGP20712A-bound structure determined here. Each profile is set to zero at its global minimum. **c,** Definition of the collective variable d used for the metadynamics simulations: the distance between the CF_3_ group of CGP20712A and the midpoint of the Cα atoms of Val/Thr^4.56^ (H4) and Val^5.45^ (H5). **d,** One-dimensional free-energy profiles along d for hβ_1_AR (light brown) and hβ_2_AR (blue), from multiple-walker well-tempered metadynamics with 16 walkers per receptor. The global minima, at d = 0.88 nm (hβ_1_AR) and 2.37 nm (β_2_AR), are labelled. **e,f,** Representative structures of CGP20712A bound to hβ_1_AR (**e**, light brown) and hβ_2_AR (**f**, blue) at d = 0.88 nm, taken as the centroid of the most populated cluster of all frames within 0.1 nm of that value. Receptors are shown as cartoons, and CGP20712A and the contacting residues as sticks; dashed lines are hydrogen bonds and fluorine contacts. Contacting residues are labelled.

To examine how CGP20712A approaches the H4-H5 insertion region, we performed multiple-walker well-tempered metadynamics (mwWT-MetaD). Simulations were initiated from inactive hβ_1_AR (PDB 7BVQ) and hβ_2_AR (PDB 2RH1) structures, in which the H4-H5 region is relatively closed compared to the CGP20712A-bound state. CGP20712A was placed fully in the extracellular solvent to avoid biasing the H4-H5 opening (Extended Data Fig. 10a). The biased collective variable (CV), d, measured the distance between the trifluoromethyl group of CGP20712A and the centre of H4 and H5 defined by two Cα atoms (Fig. 5c). Sixteen walkers were used for each receptor, providing aggregate sampling times of 14.83 μs for hβ_1_AR and 14.27 μs for hβ_2_AR. Although the profiles were well converged over most of the sampled CV range, convergence at short d remained limited. This likely reflected the slow H4 - H5 opening process and the ligand degrees of freedom. Accordingly, the one-dimensional profiles were used to compare intermediates along the binding pathway.

The global minima of the hβ_1_AR and hβ_2_AR profiles occurred at d = 0.88 and 2.37 nm, respectively (Fig. 5d). For each receptor, all frames of CVs within 0.1 nm of the corresponding minimum were clustered, and the centroid of the most populated cluster was selected as the representative structure. In hβ_1_AR, the 0.88 nm minimum positioned CGP20712A far closer to the inserted position seen in our cryo-EM structure. In the representative hβ_1_AR structure, the trifluoromethyl group points towards the H4-H5 region, forming two fluorine contacts and two hydrogen bonds (Fig. 5e). At a comparable d in hβ_2_AR, the trifluoromethyl group adopted a slightly different position as CGP20712A shifted towards the intracellular side (Fig. 5f; Extended Data Fig. 10b). Both fluorine contacts were absent, and only one hydrogen bond remained (Fig. 5f). The lower stability of the interhelical Thr^3.37^-Ser^5.46^ hydrogen bond in hβ_2_AR may facilitate this shift by creating additional space for the ligand (Extended Data Fig. 10c-d). Accordingly, the region around d = 0.9 nm was approximately 2.0 kcal/mol above the hβ_2_AR global minimum at d = 2.37 nm but defined the global minimum in hβ_1_AR (Fig. 5d). Therefore, MD simulations suggest that the H4-H5 interface is a dynamic factor in hβ_1_AR-selective ligand recognition for CGP20712A and NDD825 that insert between H4 and H5.

The MD data show that the dynamics of the H4-H5 interface make a significant contribution to the hβ_1_AR selectivity of CGP20712A. However, the dynamics of these transmembrane helices is undoubtedly affected by neighbouring helices, in addition to intracellular and extracellular loops, and of course the dynamics of these elements will be similarly affected by their neighbours. Thus, the dynamics of the H4-H5 interface will be affected by the overall dynamics of the whole receptor. A simplistic testing of the MD data through site directed mutagenesis of residues in H4 and H5 will not address global dynamics of the receptor.

## Discussion

Ligand selectivity is a key parameter to be refined during drug discovery, as off-target affinity may result in undesirable side effects. Traditionally this has been conceptualised in physical terms, such as the presence of different residues in the OBS or whether the OBS has a slightly different shape and/or size. The differences between the extended OBS in hβ_1_AR and hβ_2_AR play a significant role in the hβ_1_AR selectivity of CGP20712A, NDD825 and NDD713. In particular, His93^2.64^ in hβ_2_AR (Ile118^2.64^ in hβ_1_AR) contributes to the ligand selectivity of each of three highly selective βAR antagonists investigated here, as observed previously for some other βAR antagonists^23^. Ile94^2.65^ in hβ_2_AR (Val119^2.65^ in hβ_1_AR) also plays a lesser role in the selectivity of NDD825. However, even when all the residues in hβ_1_AR are made identical to those in hβ_2_AR, or *vice versa*, ligand selectivity cannot be converted to that observed with the other receptor. A second factor in selectivity determination is the kinetics of ligand association and dissociation. This was first shown in the βARs to explain the hβ_1_AR selectivity of noradrenaline, where the binding mode of the ligand in both hβ_1_AR and hβ_2_AR utilises identical side chains and the structures are also virtually identical^25^. We also found that this played a significant role in all the three antagonists studied here. The kinetic selectivity determined from the off-rates and on-rates were 800-1700 fold higher for hβ_1_AR over hβ_2_AR. Exchanging the extracellular vestibules of the two receptors showed that this region is necessary but not sufficient for fast association. The route to the binding site is therefore as important as the site itself.

A third factor can also influence ligand selectivity, namely subtle changes in structure. The structures of tβ_1_AR bound to CGP20712A and NDD825 showed the same unique binding mode compared to other tβ_1_AR and hβ_2_AR antagonists, with part of the ligand extending beyond the normal limits of binding of, for example catecholamines (e.g. adrenaline, noradrenaline), and extending between transmembrane helices H4 and H5. MD simulations showed that the energetics of opening this cleft were favourable for hβ_1_AR but not for hβ_2_AR. This was not intentionally designed into the ligands. Traditional medicinal chemistry and pharmacological screen (ligand affinity, as determined in pharmacological assays for both hβ_1_AR and hβ_2_AR) guided development of CGP20712A, NDD713 and NDD825 analogues with increasing selectivity to generate the final compounds published^28,29^. CGP20712A was developed over 20 years prior to the structure determination of hβ_2_AR^43^ and tβ_1_AR^16^ in 2007 and 2008, respectively. In addition, docking programmes even 10 years after the structures were determined were unable to insert either ligand correctly into the structure of tβ_1_AR, probably due to the size and complexity of the ligands^37^.

The computational tools available today for drug discovery are considerably more sophisticated and powerful, allowing more accurate docking of small molecules and new opportunities for the screening of tens of billions of molecules in a virtual library against the structure of a GPCR determined by either X-ray crystallography or, now more likely, cryo-EM^44^. However, one feature that these approaches have in common is their reliance on the (static) structure of the GPCR as it exists in the crystal or in the vitreous state frozen on cryo-EM grids. Thus, any significant structural dynamics or conformational changes that could occur upon ligand binding will not be accounted for during the computational approaches. Of course, this fact is well recognised in the development of antagonists and agonists to GPCRs, because there are abundant structures of the same receptor in different conformational states (inactive states bound to either antagonists or agonists; active states bound to full, partial or biased agonists and coupled to either a G protein, β-arrestin, G protein receptor kinase, or nothing) and comparison of the receptor structures in different states shows profound differences. For example, upon coupling to a G protein-mimetic the OBS of tβ_1_AR decreases in volume by up to 40% (depending on the ligand) due to movement of the extracellular ends of transmembrane helices H6 and H7^45^, and the conformational change is different when β-arrestin is coupled^46^. However, in the examples of CGP20712A and NDD825, the conformational state remains unchanged (inactive state), but the OBS has changed its shape upon ligand binding. Our MD simulations of ligand-free hβ_1_AR suggest that this cryptic cleft opens at a relatively low energetic cost such that it would be accessible through thermal fluctuations. In the cases of CGP20712A and NDD825, access to this cleft enhances their selectivity for hβ_1_AR over hβ_2_AR, where cleft opening is not as energetically favourable.

It is intriguing to consider whether the presence of cryptic clefts can be predicted, their role in ligand selectivity deduced and suitable ligands developed, all through computational methodologies. The computational prediction of cryptic clefts or keyholes has certainly been pursued before for hβ_1_AR^37^ and watching the thermofluctuations of a GPCR during MD simulations certainly suggests that this is not a particularly uncommon occurrence. However, utility of these observations in drug design has yet to be achieved. It is clear from these results that structure-based drug design using a structure of hβ_1_AR bound to the antagonist carazolol would never have been able to develop either CGP20712A or NDD825. What is intriguing is whether starting with a model of hβ_1_AR bound to CGP20712A would result in the selection of similar molecules or ones that have even higher selectivity for hβ_1_AR. The use of MD simulations to explore even further the extent of thermal fluctuations at the H4-H5 interface may suggest that starting with a model with a wider cryptic cleft may be even more beneficial. There may be considerable potential for applying similar considerations in drug design to, for example, the serotonin receptor family which also faces considerable challenges in achieving subtype selectivity. We anticipate that a better understanding of the structural and energetic mechanisms of hβ_1_AR selective ligands may inform future approaches for the design of highly selective GPCR ligands.

## Methods

### Chemistry methodology

Chemicals and solvents were purchased from standard suppliers and used without further purification. Merck Kieselgel 60, 230-400 mesh, for flash column chromatography (FCC) was supplied by Merck KgaA (Darmstadt, Germany) and deuterated solvents were purchased from Goss International Limited (England) and Sigma-Aldrich Company Ltd (England).

Unless otherwise stated, reactions were carried out at ambient temperature. Reactions using a microwave reactor were carried out in a CEM Discover system, with an Explorer 48 autosampler. Reactions were monitored by thin layer chromatography on commercially available precoated aluminium-backed plates (Merck Kieselgel 60 F_254_). Visualisation was by examination under UV light (254 and 366 nm). General staining carried out with KMnO_4_ or phosphomolybdic acid. A solution of Ninhydrin (in ethanol) was used to visualize primary and secondary amines. All organic extracts collected after aqueous work-up procedures were dried over anhydrous MgSO_4_ or Na_2_SO_4_ before gravity filtering and evaporation to dryness. Organic solvents were evaporated *in vacuo* at 40°C (water bath temperature). Purification using preparative layer chromatography (PLC) was carried out using Fluka silica gel 60 PF_254_ containing gypsum (200 mm x 200 mm x 1 mm). Flash chromatography was performed using Merck Kieselgel 60 (0.040-0.063 mm).

Melting points (Mp) were recorded on a Reichert 7905 apparatus or Perkin Elmer Pyris 1 differential scanning calorimeter and were uncorrected. Fourier transform – infrared (FT-IR) spectra were recorded as thin films or KBr discs in the range of 4000 – 500 cm^-1^ using an Avatar 360 Nicolet FT-IR spectrophotometer. High resolution mass spectra (HRMS) – time of flight, electrospray (TOF ES +/-) were recorded on a Waters 2795 separation module/micromass LCT platform. ^1^H NMR spectra were recorded on a Bruker-AV 400 at 400.13 MHz. ^13^C NMR spectra were recorded at 101.62 MHz. Chemical shifts (δ) are recorded in parts per million (ppm) with reference to the chemical shift of the deuterated solvent/an internal tetramethylsilane (TMS) standard. Coupling constants (*J*) and carbon-fluorine coupling constants (*J_CF_*) are recorded in Hz and the significant multiplicities described by singlet (s), doublet (d), triplet (t), quadruplet (q), broad (br), multiplet (m), doublet of doublets (dd), doublet of triplets (dt). Spectra were assigned using appropriate COSY, distortionless enhanced polarisation transfer (DEPT), HSQC and HMBC sequences. Unless otherwise stated all spectra were recorded in CDCl_3_.

Analytical reverse-phase high-performance liquid chromatography (RP-HPLC) was performed on a Shimadzu UFLCXR system coupled to an Applied Biosystems API2000 using system 1 All retention times (*t*_R_) are quoted in minutes.

System 1: Gemini-NX 3u-110A, 50×2mm column thermostated at 40°C, a flow rate of 0.5 mL/min and UV detection at 220 and 254nm. Pre-equilibration run for one minute at 10% solvent B, then 10% - 98% solvent B over 2 minutes, then 98% solvent B for 2 minutes, then 98% - 10% solvent B in 0.5 minutes then 10% for one minute. Solvent A: 0.1% FA in water; solvent B: 0.1% FA in MeCN.

Preparative HPLC was performed using a Phenomenex Onyx Monolithic reverse phase C_18_ column (100 x 10 mm), a flow rate of 14.10 mL/min and UV detection at 287 nm. Samples were run in 5% - 95% solvent B over 10 minutes. Solvent A: 0.1% FA in water; solvent B: 0.1% FA in MeCN.

#### General Method A: Ring-opening of (*R*)-2-((cyclopropylmethoxy)methyl)-6-(((*S*)-oxiran-2-yl)methoxy)-2,3-dihydrobenzo[*b*][1,4]dioxine (**19**) with amine hydrochloride salts

(*R*)-2-((Cyclopropylmethoxy)methyl)-6-(((*S*)-oxiran-2-yl)methoxy)-2,3-dihydrobenzo[*b*][1,4]dioxine (**19**) (100 mg) and amine (1.5 eq) were weighed into a 10 mL MW reactor vial with a micro-stirrer bar. DIPEA (1.6 eq) was added, before adding propan-2-ol/MeCN/H_2_O (7:2:1, 5 mL). The mixture was sonicated, and the resulting solution or suspension heated at 90 °C in the MW reactor (dynamic program, with maximum pressure = 200 psi, maximum power = 300W) with stirring for 60 minutes. The reaction mixture was then concentrated under reduced pressure, and the crude residue further purified by flash column chromatography (FCC) (eluent 1N NH_3_ in MeOH/dichloromethane with a gradient from 1:99 to a maximum of 20:80).

**Scheme 1:**
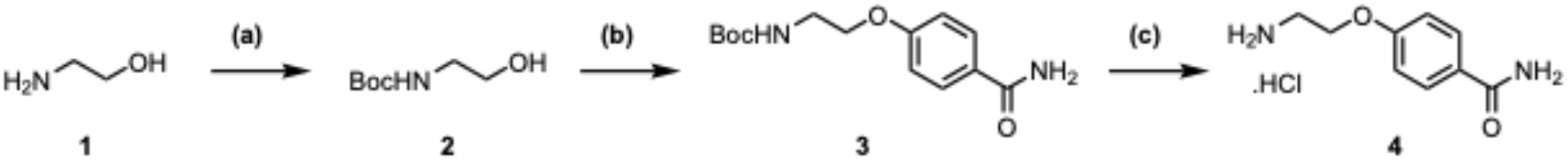
Synthesis of 4-(2-aminoethoxy)benzamide hydrochloride (3)*^a^* *^a^ Reagents and conditions.* (a) Di-*tert*-butyl dicarbonate (Boc_2_O), water, tetrahydrofuran (THF), 99%; (b) 4-hydroxybenzamide, PPh_3_, diisopropylazodicarboxylate (DIAD), THF, 37%; (c) 4M HCl/1,4-dioxane, 100%.

*tert-Butyl 2-hydroxyethylcarbamate (**2**).* Ethanolamine (**1**) (4.940 mL, 81.86 mmol) was dissolved in water (75 mL) and THF (35 mL). To this was added Boc_2_O (19.650 g, 90.05 mmol, 1.1 eq) and the mixture stirred at room temperature overnight. After removal of THF *in vacuo*, the aqueous slurry was extracted with dichloromethane (DCM) (3 x 30 mL). The organic layer was concentrated to give 13.077 g (99%) of clear colourless oil employed directly in the following step. FT-IR: 3360, 2978, 2934, 1689, 1527, 1393, 1367, 1252, 1173; ^1^H NMR δ 5.18 (br s, 1H), 3.64 (dt, *J* = 5.5/4.9 Hz, 2H), 3.26 – 3.35 (br m, 1H), 3.23 (dt, *J* = 5.1/5.1 Hz, 2H), 1.40 (s, 9H); ^13^C NMR δ 156.9, 79.7, 62.3, 43.2, 28.5; *m/z* HRMS (TOF ES^+^) C_7_H_16_NO_3_ [MH]^+^ calcd 162.1125; found 162.1129.

*tert-Butyl 2-(4-carbamoylphenoxy)ethylcarbamate (**3**). tert*-Butyl 2-hydroxyethylcarbamate (**2**) (5.343 g, 33.14 mmol), PPh_3_ (10.431 g, 39.77 mmol, 1.2 eq) and 4-hydroxybenzamide (5.00 g, 36.46 mmol, 1.1 eq) were dispersed in THF (100 mL), before addition of DIAD (7.83 mL, 39.77 mmol, 1.2 eq) as a solution in THF (20 mL), dropwise. After addition was complete, the mixture was stirred at room temperature for 2 days. Over this time a precipitate formed to give a white suspension which was vacuum filtered, and washed with Et_2_O to afford the title compound 3.47 g (37%) as a white solid. ^1^H NMR (DMSO-*d_6_*) δ 7.83 (d, *J* = 8.4 Hz, 2H), 7.17 (br s, 1H), 7.02 (br s, 1H), 6.96 (d, *J* = 8.4 Hz, 2H), 3.95 – 4.10 (m, 2H), 3.18 – 3.44 (m, 2H), 1.38 (s, 9H); ^13^C NMR (DMSO-*d_6_*) δ 167.4, 160.8, 155.7, 129.4, 126.6, 113.9, 77.8, 66.5, 28.2, 21.9; *m/z* MS (TOF ES^+^) C_14_H_21_N_2_O_4_ [MH]^+^ calcd 281.1; found 281.2; RP-HPLC *t*_R_: 2.34 min (System 1).

*4-(2-Aminoethoxy)benzamide hydrochloride (**4**). tert*-Butyl 2-(4-carbamoylphenoxy)ethylcarbamate (**3**) (3.456 g, 12.33 mmol) was dissolved in DCM (50 mL), with addition of MeOH (10 mL) to aid dissolution. 4M HCl/1,4-dioxane (60 mL) was added, and the mixture stirred at room temperature for 3 hours. The reaction mixture was flooded with petroleum ether 40-60, and the resulting precipitate collected by vacuum filtration, before drying to give the title compound. Yield: 100%, white solid; ^1^H NMR (DMSO-*d_6_*) δ 11.29 (br s, 1H), 8.41 (br s, 3H), 7.87 (d, *J* = 8.9 Hz, 2H), 7.01 (d, *J* = 8.9 Hz, 2H), 4.25 (t, *J* = 5.2 Hz, 2H), 3.15 – 3.25 (m, 2H); ^13^C NMR (DMSO-*d_6_*) δ 167.3, 160.2, 129.4, 127.1, 114.1, 64.4, 38.2.

**Scheme 2:**
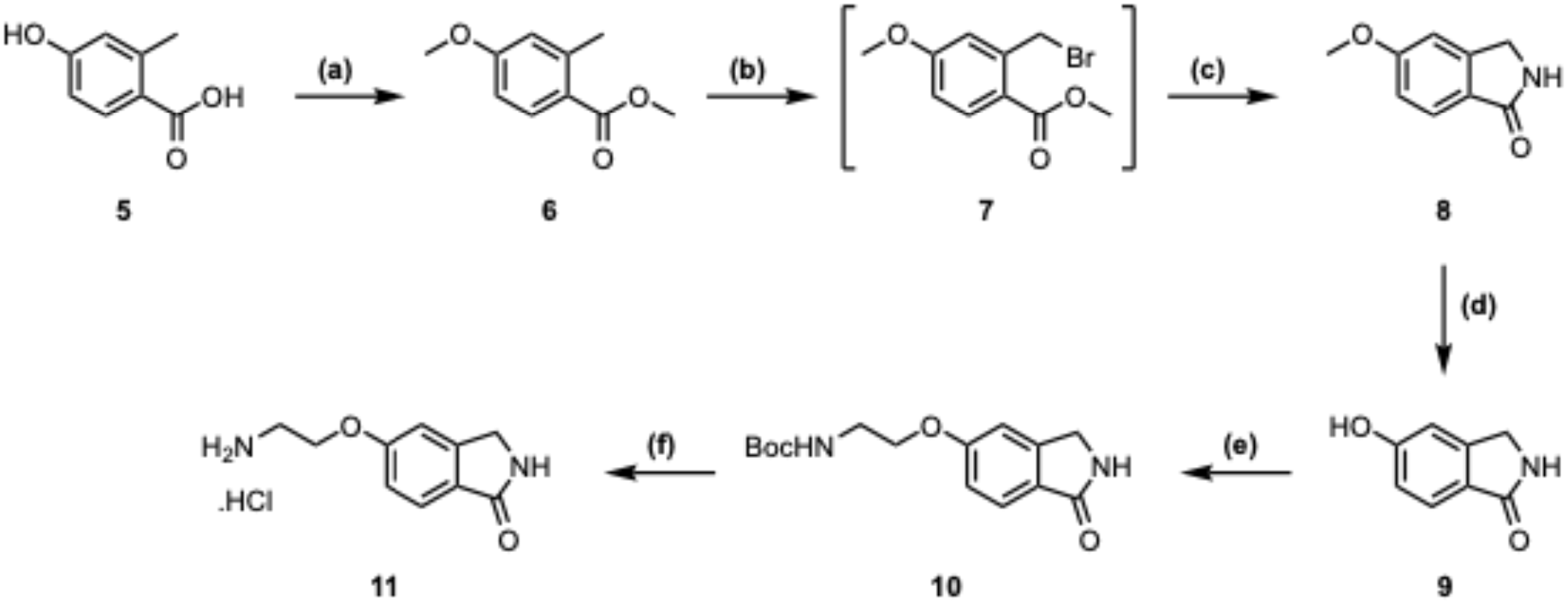
Synthesis of 5-(2-aminoethoxy)isoindolin-1-one hydrochloride (11)*^a^* *^a^ Reagents and conditions.* (a) K_2_CO_3_, MeI, MeCN, reflux, 100%; (b) *N*-bromosuccinimide (NBS), 1,1′-azobis(cyclohexanecarbonitrile) (ABCN), CCl_4_, reflux; (c) 7M NH_3_/MeOH, 70 °C, 63% (2 steps); (d) 1M BBr_3_/DCM, 0 °C to rt, 100%; (e) *tert*-butyl 2-bromoethylcarbamate, Cs_2_CO_3_, DMF, rt to 50 °C, 100%; (f) 4M HCl/1,4-dioxane, rt, 100%.

*Methyl 4-methoxy-2-methylbenzoate (**6**).* 4-Hydroxy-2-methylbenzoic acid (**5**) (21.0 g, 138 mmol), K_2_CO_3_ (47.7 g, 345 mmol, 2.5 eq) and MeI (18.9 mL, 304 mmol, 2.2 eq) were dispersed in MeCN (400 mL) and heated under reflux for 6.5 hours. TLC analysis (EtOAc/PE 1:1) indicated the reaction was complete. The mixture was cooled, and filtered, before concentrating the filtrate. The residue was dispersed in water (200 mL), before extracting with DCM (3 x 100 mL). The combined organic layers were concentrated to give 25.0 g (100%) of the title compound as a yellow oil. ^1^H NMR δ 7.93 (d, *J* = 9.4 Hz, 2H), 6.69 – 6.80 (m, 2H), 3.85 (s, 3H), 3.83 (s, 3H), 2.60 (s, 3H); ^13^C NMR δ 167.7, 162.4, 143.3, 133.1, 121.9, 117.1, 111.1, 55.4, 51.7, 22.5.

*5-Methoxyisoindolin-1-one (**8**).* Methyl 4-methoxy-2-methylbenzoate (**6**) (34.286 g, 190.27 mmol) was dissolved in CCl_4_ (500 mL), to this was added NBS (37.249 g, 209.29 mmol, 1.1 eq), followed by ABCN (9.298 g, 38.05 mmol, 0.2 eq). The resulting pale-yellow suspension was heated under reflux under an atmosphere of nitrogen overnight. The mixture was cooled, and the precipitate filtered and washed with CCl_4_. The filtrate was washed with 10% Na_2_S_2_O_3 (aq)_ (200 mL), before concentrating to give 60.3 g of yellow oil. This was reacted without further purification.

The crude methyl 2-(bromomethyl)-4-methoxybenzoate **(7)** (59.8 g, 230.8 mmol) was heated in 7M NH_3_ in MeOH (500 mL) at 70°C for 3.75 hours. TLC analysis (EtOAc/petroleum ether 40-60 1:1 and MeOH/DCM 1:9) indicated starting material had been consumed. The mixture was cooled, before concentrating. The yellow residue was dispersed in water (200 mL) and Et_2_O (200 mL). The resulting mixture was shaken before filtering the resulting precipitate. This was washed with Et_2_O, and afforded 16.677 g of the title compound as a yellow solid after drying.

TLC analysis of the aqueous layer, Et_2_O layer and precipitate against a previously purified and characterized sample of 5-methoxyisoindolin-1-one (**8**), indicated the precipitate was pure product, the Et_2_O layer contained mainly impurities with some product and the remaining product was trapped in the aqueous layer. The Et_2_O layer was combined with ethereal precipitate washings and extracted with water (80 mL), causing selective extraction of the desired compound. The combined Et_2_O layers were then concentrated to around 80 mL in volume, before further extraction with water (2 x 100 mL). The combined aqueous layers were then saturated with NaCl, before extracting with EtOAc (2 x 100 mL) and 10% MeOH/EtOAc (2 x 100 mL). The combined EtOAc containing extracts were dried and concentrated, and the resulting residue sonicated in Et_2_O. The resulting yellow precipitate was collected by filtration and washed with petroleum ether 40-60. Combined weight of collected solid = 19.544 g (63% over 2 steps). ^1^H NMR (DMSO-*d_6_*) δ 8.30 (br s, 1H), 7.56 (d, *J* = 8.4 Hz, 1H), 7.12 (d, *J* = 1.8 Hz, 1H), 7.01 (dd, *J* = 8.4/2.3 Hz, 1H), 4.31 (s, 2H), 3.82 (s, 3H); ^13^C NMR (DMSO-*d_6_*) δ 169.8, 162.2, 146.5, 125.1, 124.0, 114.7, 108.1, 55.6, 44.7.

*5-Hydroxyisoindolin-1-one (**9**).* 5-Methoxyisoindolin-1-one (**8**) (2.867 g, 17.57 mmol) was dispersed in DCM (20 mL) over an ice bath and stirred for 20 minutes, before dropwise addition of 1M BBr_3_/DCM (53 mL, 53 mmol, 3 eq). This resulted in formation of a brown solution. After addition was complete, the mixture was warmed to room temperature, and stirred for 2 hours, causing an off-white precipitate to form. TLC analysis (MeOH/DCM 1:9) indicated the reaction was complete. The mixture was cooled over an ice bath, before careful quenching with MeOH (10 mL). After returning to room temperature, the mixture was concentrated under reduced pressure. The resulting off-white solid was dispersed in water (100 mL) and stirred at room temperature overnight, before heating at 60°C for 1 hour. The suspension was then cooled and filtered, before drying to give 2.657 g (100%) of the title compound as an off-white solid. ^1^H NMR (DMSO-*d_6_*) δ 10.05 (s, 1H), 8.16 (br s, 1H), 7.46 (d, *J* = 8.2 Hz, 1H), 6.86 – 6.92 (m, 1H), 6.83 (dd, *J* = 8.2/2.0 Hz, 1H), 4.25 (s, 2H); ^13^C NMR (DMSO-*d_6_*) δ 170.1, 160.6, 146.6, 124.2, 123.6, 115.4, 109.7, 44.6.

*tert-Butyl 2-(1-oxoisoindolin-5-yloxy)ethylcarbamate (**10**).* 5-Hydroxyisoindolin-1-one (**9**) (16.0 g, 107.27 mmol) was dissolved in DMF (50 mL), before adding Cs_2_CO_3_ (38.45 g, 118.0 mmol, 1.1 eq) and stirred at room temperature for 1 hour. A solution of *tert*-butyl 2-bromoethylcarbamate (26.44 g, 118.0 mmol, 1.1 eq) in DMF (150 mL) was added dropwise, before allowing the mixture to stir at room temperature for 2 days. Cs_2_CO_3_ (10.48 g, 0.3 eq) and *tert*-butyl 2-bromoethylcarbamate (7.21 g, 0.3 eq) were added, and stirring continued for 4 hours, before increasing the temperature to 50 °C for 4 hours and then stirred at room temperature overnight. The mixture was concentrated under reduced pressure, before adding EtOAc (500 mL) and washing with 2M NaOH _(aq)_ (400 mL) and brine (400 mL), allowing any emulsion to partition into the aqueous layer. After separation, Tthe organic layer was then flooded with petroleum ether 40-60, causing precipitation of the product. This was collected by filtration (vacuum),and washed with petroleum ether 40-60. The aqueous layers were combined and saturated with NaCl, before being extracted with EtOAc (2 x 300 mL). The organic extracts were then combined with the organic filtrate and concentrated… The concentrated organic layer was dispersed in a small amount of EtOAc with heating, before adding petroleum ether 40-60 and vacuum filtering the precipitate. TLC analyses of the precipitate and filtrate showed complete isolation of the desired compound. Combined yield = 31.9 g (100%) as an off-white solid. ^1^H NMR δ 7.77 (d, *J* = 8.4 Hz, 1H), 6.99 (dd, *J* = 8.4/2.1 Hz, 1H), 6.91 – 6.95 (m, 1H), 6.48 (br s, 1H), 5.00 (br s, 1H), 4.40 (s, 2H), 4.08 (t, *J* = 5.1 Hz, 2H), 3.50 – 3.62 (m, 2H), 1.45 (s, 9H); ^13^C NMR δ 167.6, 161.4, 156.0, 143.3, 133.1, 122.2, 117.5, 111.5, 79.8, 67.3, 51.7, 40.2, 28.5.

*5-(2-Aminoethoxy)isoindolin-1-one hydrochloride (**11**). tert*-Butyl 2-(1-oxoisoindolin-5-yloxy)ethylcarbamate (**10**) (2.0 g, 6.84 mmol) was dissolved in DCM (20 mL). 4M HCl/1,4-dioxane (30 mL) was added, and the mixture stirred at room temperature for 3 hours. The reaction mixture was flooded with petroleum ether 40-60, and the resulting precipitate collected by vacuum filtration, before drying to give the title compound. Yield: 100%, white solid; ^1^H NMR (DMSO-*d_6_*) δ 8.20 – 8.45 (m, 4H), 7.59 (d, *J* = 8.4 Hz, 1H), 7.16 (d, *J* = 1.8 Hz, 1H), 7.06 (dd, *J* = 8.4/2.2 Hz, 1H), 4.32 (s, 2H), 4.27 (t, *J* = 5.1 Hz, 2H), 3.17 – 3.28 (m, 2H); ^13^C NMR (DMSO-*d_6_*) δ 169.7, 160.7, 146.5, 125.7, 124.1, 115.4, 108.8, 64.7, 44.8, 38.2.

**Scheme 3:**
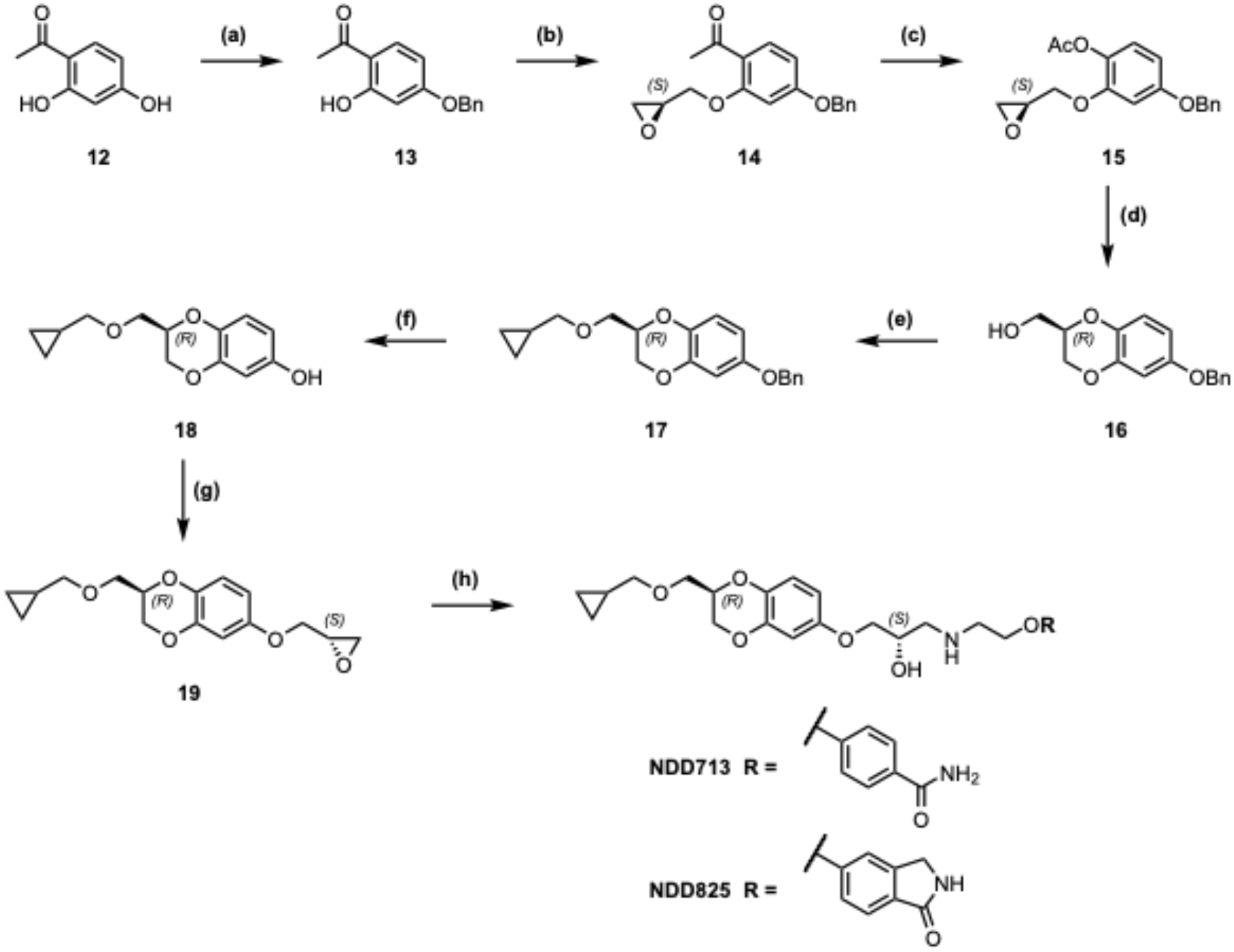
Synthesis of NDD713 and NDD825*^a^* *^a^ Reagents and conditions.* (a) K_2_CO_3_, MeCN, reflux, 30 min, then BnBr, reflux, 3 h, 95%; (b) i. NaH, anhydrous *N*,*N*-dimethylformamide (DMF), rt, 30 min; ii. *S*-glycidyl 3-nitrobenzenesulfonate, 60 °C, 5 h 89%; (c) *meta*-chloroperbenzoic acid (*m*-CPBA), NaHCO_3_, DCM, 40 °C, 12 h, 98%; (d) aq NaOH, THF, 16 h, rt, 92%; (e) cyclopropylmethyl bromide, anhydrous *N-N-*dimethylacetamide (DMA), rt, 10 min, then NaO*^t^*Bu, −5 °C to rt, 16 h, 94%; (f) 10% Pd(C), DCM/MeOH (8:2), H_2_, rt, 12 h, 100%; (g) i. NaH, anhydrous DMF, rt, 30 min; ii. *S*-glycidyl 3-nitrobenzenesulfonate, 60°C, 12 h, 95%;

*4-Benzyloxy-2-hydroxyacetophenone (**13**).* 2,4-Dihydroxyacetophenone (85.0 g, 558.67 mmol) was dissolved in MeCN (700 mL) and K_2_CO_3_ (92.653 g, 670.40 mmol, 1.2 eq) was added. The mixture was heated under reflux for 30 min, giving a peach-coloured suspension. BnBr (73.1 mL, 614.53 mmol, 1.1 eq.) as a solution in MeCN (200 mL) was added dropwise to the reaction flask over 40 min, and the mixture was then heated under reflux for a further 2-3 h. TLC analysis (EtOAc/petroleum ether 40-60, 3:7) indicated complete disappearance of the starting material. The mixture was cooled and inorganic solids were separated by filtration. The organic layer was concentrated to a slurry, and was then dispersed in aq 1 M HCl (300 mL) and extracted with DCM (3 x 400 mL). The combined organic layers were concentrated to give a beige solid. The crude product was recrystallized by dissolution in a minimum amount of EtOAc (heating was required) and then careful addition of petroleum ether until crystal deposition started. Off-white crystals were finally separated by gravity filtration and dried (128 g, 95%). ^1^H NMR (CDCl_3_) δ 12.73 (s, 1H), 7.66 - 7.62 (m, 1H), 7.44 - 7.33 (m, 5H), 6.53 - 6.50 (m, 1H+1H), 5.10 (s, 2H), 2.56 (s, 3H); ^13^C NMR (CDCl_3_) 202.7, 165.3, 136.0, 132.5, 128.8, 128.5, 127.7, 114.2, 108.3, 102.0, 70.3, 26.4.

*1-(4-(Benzyloxy)-2-(((S)-oxiran-2-yl)methoxy)phenyl)ethan-1-one (**14**).* To a stirred solution of NaH (476.74 mmol, 18.267 g), in anhydrous DMF (250 mL) under a N_2_ atmosphere, was added dropwise a solution of 4-benzyloxy-2-hydroxy-1-acetophenone (**13**) (433.40 mmol, 105.0 g) in anhydrous DMF (600 mL). The mixture was allowed to stir at room temperature for 30 min, then (*S*)-glycidyl-3-nitrobenzenesulfonate (433.40 mmol, 112.354 g) in anhydrous DMF (350 mL) was added dropwise at room temperature. The mixture was slowly heated to 60 °C and stirred for 48 h. Completion of the reaction was monitored by TLC and LC-MS. After 48 h the reaction mixture was quenched with saturated aq NH_4_Cl solution (*ca.* 50 mL) and then with H_2_O (100 mL). Solvent was removed under high vacuum and the residue was suspended in H_2_O (350 mL). The aqueous phase was extracted with EtOAc (3 x 500 mL). The combined organic layers were successively washed with 1 M aq NaOH (3 x 300mL), H_2_O (2 x 300 mL), brine (1 x 400mL), finally dried over Na_2_SO_4_ and filtered. The organic phase was evaporated to give a dark brown solid. The crude product was purified by column chromatography (eluent petroleum ether 40-60/EtOAc, 7:3) to afford the title compound as an off-white solid (105 g, 89%). Unreacted 4-benzyloxy-2-hydroxy-1-acetophenone (**13**) (9.0 g) was also recovered. ^1^H NMR (CDCl_3_) δ 7.83 (d, *J* = 8.8 Hz, 1H), 7.43 - 7.32 (m, 5H), 6.62 (dd, *J* = 8.6/2.3 Hz, 1H), 6.52 (d, *J* = 2.4 Hz, 1H), 5.10 (s, 2H), 4.32 (dd, *J* = 10.9/2.9 Hz, 1H), 3.95 (dd, *J* = 10.9/6.0 Hz, 1H), 3.41 - 3.37 (m,1H), 2.92 (dd, *J* = 4.8/4.3 Hz, 1H), 2.76 (dd, *J* = 4.9/2.8 Hz, 1H), 2.62 (s, 3H). ^13^C NMR (CDCl_3_) 197.6, 163.6, 159.8, 136.2, 132.9, 128.8, 128.4, 127.7, 121.7, 106.7, 100.3, 70.4, 69.7, 49.9, 44.7, 32.1.

*4-(Benzyloxy)-2-(((S)-oxiran-2-yl)methoxy)phenyl acetate (**15**).* To a stirred solution of 1-(4- (benzyloxy)-2-(((*S*)-oxiran-2-yl)methoxy)phenyl)ethan-1-one (**14**) (103.0 g, 345.25 mmol) in DCM (1 L) was added *m*-CPBA (137.031 g, 555.855 mmol) and solid NaHCO_3_ (203.03 g, 2416.76 mmol). The suspension was stirred at 40 °C (reflux) overnight (12 h). Completion of the reaction was checked by LC-MS. To the cooled suspension was added 10% aq Na_2_S_2_O_3_ solution (1 L) and the mixture was stirred for 15 min. Inorganic salts were removed by filtration.

The organic layer was washed sequentially with 10% aq Na_2_S_2_O_3_ solution (2 x 300 mL), saturated aq NaHCO_3_ solution (2 x 500mL), H_2_O (1 x 600 mL), and finally with brine (1 x 600 mL). After drying over Na_2_SO_4_ and filtration, solvent was removed by rotary evaporation to afford the title product as a crude thick orange oily, requiring no further purification (106.5 g, 98%). ^1^H NMR (CDCl_3_) δ 7.49 - 7.29 (m,5H), 6.95 (d, *J* = 8.7 Hz, 1H), 6.64 (d, *J* = 2.8 Hz, 1H), 6.55 (dd, *J* = 8.7/2.8 Hz, 1H), 5.03 (s, 2H), 4.20 (dd, *J* = 11.2/3.0 Hz, 1H), 3.95 (dd, *J* = 11.2/5.5 Hz, 1H), 3.32 - 3.28 (m, 1H), 2.87 (dd, *J* = 4.9/4.2 Hz, 1H), 2.71 (dd, *J* = 4.9/2.7 Hz, 1H), 2.30 (s, 3H); ^13^C NMR (CDCl_3_) 169.6, 157.6, 150.8, 136.8, 134.3, 128.8, 128.2, 127.7, 123.1, 106.2, 102.7, 77.4, 70.6, 69.7, 50.1, 44.6, 20.7.

*((R)-6-(Benzyloxy)-2,3-dihydrobenzo[b][1,4]dioxin-2-yl)methanol (**16**).* To a stirred solution of 4-(benzyloxy)-2-(((*S*)-oxiran-2-yl)methoxy)phenyl acetate (**15**) (337.22 mmol, 106 g) in THF (700 mL) was added aq 2 M NaOH solution (202.33 mL, 404.67 mmol). The mixture was stirred at room temperature for 18 h. Completion of the reaction was monitored by TLC/LC-MS. When completed, THF was removed in vacuo and the residue was redissolved in H_2_O (200 mL). The solution was extracted with EtOAc (3 x 600 mL) and the combined organic extracts were washed with H_2_O (1 x 600 mL) and brine (1 x 600 mL), dried over Na_2_SO_4_, filtered and solvent was removed *in vacuo* to afford a transparent thick oil which turned into a white solid on drying, requiring no further purification (85.2 g, 92%). ^1^H NMR (CDCl_3_) δ 7.44 - 7.30 (m,5H), 6.81 (d, *J* = 8.9 Hz, 1H), 6.56 (d, *J* = 3.0 Hz, 1H), 6.52 (dd, *J* = 8.8/2.9 Hz, 1H), 4.99 (s, 2H), 4.27 (dd, *J* = 11.2/2.2 Hz, 1H), 4.22 - 4.16 (m, 1H), 4.08 (dd, *J* = 11.2/7.7 Hz, 1H), 3.90 - 3.78 (m,2H), 2.27 (br s, 1H). ^13^C NMR (CDCl_3_) 153.6, 143.4, 137.3, 137.2, 128.7, 128.0, 127.6, 117.5, 108.6, 103.9, 73.3, 70.7, 65.5, 61.8.

*(R)-6-(Benzyloxy)-2-((cyclopropylmethoxy)methyl)-2,3-dihydrobenzo[b][1,4]dioxine (**17**).* ((*R*)- 6-(Benzyloxy)-2,3-dihydrobenzo[*b*][1,4]dioxin-2-yl)methanol (**16**) (15 g, 55.1 mmol) was weighed into a dry flask under an atmosphere of N_2_. To this was added cyclopropylmethyl bromide (6.411 mL, 66.1 mmol, 1.2 eq) and dry DMA (250 mL). The mixture was stirred at room temperature for 10 min before cooling to −5 °C in an ice/NaCl bath. NaO^t^Bu (7.412 g, 77.12 mmol, 1.4 eq) was added slowly portion-wise, before leaving the reaction stirring in the ice bath overnight, allowing it to reach room temperature. TLC analysis (eluent EtOAc/petroleum ether 40-60 3:7) indicated presence of some starting material. Additional cyclopropylmethyl bromide (1 mL) and NaO^t^Bu (1.5 g) were added portion-wise and stirred for 3 h. TLC revealed complete conversion. The reaction was quenched with H_2_O (50 mL) before concentrating. H_2_O (120 mL) was added before extracting with Et_2_O (3 x 120 mL). The combined organic phases were washed with brine (250 mL) before concentrating to give the desired product as a colourless viscous oil, which became a white semi solid on further drying (16.882 g, 94%). This compound was taken through to the next step without further purification.

^1^H NMR (CDCl_3_) δ 7.43 - 7.29 (m,5H), 6.81 (d, *J* = 8.6 Hz, 1H), 6.54 (d, *J* = 2.9 Hz, 1H), 6.50 (dd, *J* = 8.7/3.0 Hz, 1H), 4.98 (s, 2H), 4.32 - 4.26 (m, 2H), 4.10 - 4.04 (m, 1H), 3.76 - 3.72 (m,1H), 3.66 - 3.61 (m, 1H), 3.41 - 3.33 (m, 2H), 1.12 - 1.02 (m, 1H), 0.61 - 0.49 (m, 2H), 0.28 - 0.17 (m, 2H); ^13^C NMR (CDCl_3_) 153.4, 143.4, 137.4, 137.3, 128.5, 127.9, 127.5, 117.5, 108.3, 103.7, 76.4, 71.9, 70.5, 69.0, 65.9, 10.5, 3.1, 3.0. *m/z* MS (TOF ES^+^) C_20_H_25_O_4_ [MH]^+^ calcd 327.2; found 327.3; RP-HPLC *t*_R_: 3.22 min (System 1).

*(R)-2-((Cyclopropylmethoxy)methyl)-2,3-dihydrobenzo[b][1,4]dioxin-6-ol (**18**).* To a solution of (*R*)-6-(Benzyloxy)-2-((cyclopropylmethoxy)methyl)-2,3-dihydrobenzo[*b*][1,4]dioxine (**17**) (51.47 mmol, 16.8 g) in DCM/MeOH (8:2; 200mL) was added 10% Pd/C (1.7 g, 0.1 eq by weight) in water (4 mL) (to avoid a very exothermic reaction). The suspension was degassed with cycles of vacuum and N_2_ flushes, before placing the suspension under an atmosphere of H_2_. The suspension was stirred at room temperature for 18 h. Completion of the reaction was monitored by LC-MS/TLC. The mixture was then filtered through a bed of celite with DCM (100 mL) washing. The combined organic layers were concentrated to leave an oil with some water. To this was added DCM (200 mL) and the solution was dried over Na_2_SO_4_. DCM was rotary evaporated to obtained a brown oil, which was filtered through a silica plug (petroleum ether 40-60/EtOAc 7:3) to give a colourless gum (12.1 g, quantitative). ^1^H NMR (CDCl_3_) δ 6.72 (d, *J* = 8.8 Hz, 1H), 6.40 (d, *J* = 2.9 Hz, 1H), 6.33 (dd, *J* = 8.7/3.0 Hz, 1H), 5.56 (br s, 1H), 4.30 - 4.25 (m, 2H), 4.04 (dd, *J* = 11.5/7.9 Hz, 1H), 3.73 (app dd, *J* = 10.6/5.2 Hz, 1H), 3.64 (app dd, *J* = 10.4/5.4 Hz, 1H), 3.37 (app d, *J* = 6.6 Hz, 2H), 1.11 - 1.01 (m, 1H), 0.60 - 0.48 (m, 2H), 0.26 - 0.15 (m, 2H); ^13^C NMR (CDCl_3_) 150.2, 143.5, 137.0, 117.7, 108.6, 104.2, 76.7, 71.9, 69.1, 66.0, 10.5, 3.2, 3.1. RP-HPLC *t*_R_: 2.58 min (System 1)

*(R)-2-((Cyclopropylmethoxy)methyl)-6-(((S)-oxiran-2-yl)methoxy)-2,3-dihydrobenzo[b][1,4]dioxine (**19**).* To a stirred suspension of NaH (56.57 mmol, 2.168 g, 1.1 eq) in anhydrous DMF (30 mL) under N_2_ was added dropwise a solution of (*R*)-2- ((cyclopropylmethoxy)methyl)-2,3-dihydrobenzo[*b*][1,4]dioxin-6-ol (**18**) (51.43 mmol, 12.05 g, 1 eq) in anhydrous DMF (60 mL). The mixture was allowed to stir at room temperature for 30 min and then a solution of (*S*)-glycidyl-3-nitrobenzenesulfonate (51.43 mmol, 13.333 g, 1 eq) in anhydrous DMF (30 mL) was added dropwise under N_2_ at room temperature. The mixture was slowly heated to 60 °C and stirred for 36 h. The completion of the reaction was monitored by TLC. The DMF solvent was evaporated under high vacuum and the residue quenched with water (50 mL). The crude product was extracted with EtOAc (3 x 60 mL). The combined organic layers were successively washed with 1 M aq NaOH (2 x 100 mL), H_2_O (2 x 100 mL) and with brine (1 x 100 mL). After drying over Na_2_SO_4_ and filtration, EtOAc was removed *in vacuo* to give a crude product as a light orange oil of sufficient purity to carry forward (14.2 g, 95%) to the next step. ^1^H NMR (CDCl_3_) δ 6.78 (d, *J* = 8.9 Hz, 1H), 6.47 (d, *J* = 2.8 Hz, 1H), 6.43 (dd, *J* = 8.8/2.9 Hz, 1H), 4.33 - 4.23 (m, 2H), 4.12 (dd, *J* = 11.1/3.2 Hz, 1H), 4.07 - 4.02 (m, 1H), 3.87 (dd, *J* = 11.0/5.7 Hz, 1H), 3.72 (app dd, *J* = 10.5/5.2 Hz, 1H), 3.62 (app dd, *J* = 10.4/5.9 Hz, 1H), 3.39 - 3.33 (m, 2H), 3.32 - 3.28 (m, 1H), 2.87 (dd, *J* = 5.0/4.2 Hz, 1H), 2.72 (dd, *J* = 5.0/2.8 Hz, 1H), 1.11 - 1.01 (m, 1H), 0.60 - 0.48 (m, 2H), 0.26 - 0.15 (m, 2H); ^13^C NMR (CDCl_3_) 153.1, 143.5, 137.7, 117.6, 108.2, 103.7, 76.6, 72.0, 69.5, 69.3, 69.1, 66.1, 50.3, 44.8, 10.5, 3.14, 3.05; RP-HPLC *t*_R_: 2.87 min (System 1)

*4-(2-((S)-3-((R)-2-((Cyclopropylmethoxy)methyl)-2,3-dihydrobenzo[b][1,4]dioxin-6-yloxy)-2-hydroxypropylamino)ethoxy)benzamide (**NDD713**).* Prepared by general method A, using 4- (2-aminoethoxy)benzamide hydrochloride (**4**). Yield: 29%, white solid; Mp = 136 – 138 °C; ^1^H NMR (DMSO-*d_6_*) δ 7.83 (d, *J* = 8.8 Hz, 2H), 7.82 (br s, 1H), 7.16 (br s, 1H), 6.97 (d, *J* = 8.8 Hz, 2H), 6.77 (d, *J* = 8.8 Hz, 1H), 6.46 (d, *J* = 2.8 Hz, 1H), 6.42 (dd, *J* = 8.8/2.9 Hz, 1H), 4.98 (br d, *J* = 4.0 Hz, 1H), 4.28 (dd, *J* = 11.3/2.3 Hz, 1H), 4.19 – 4.27 (m, 1H), 4.07 (t, *J* = 5.5 Hz, 2H), 3.96 (dd, *J* = 11.3/7.3 Hz, 1H), 3.74 – 3.89 (m, 3H), 3.56 – 3.66 (m, 2H), 3.29 (d, *J* = 6.8 Hz, 2H), 2.91 (t, *J* = 5.5 Hz, 2H), 2.72 (dd, *J* = 11.8/3.9 Hz, 1H), 2.62 (dd, *J* = 11.8/6.4 Hz, 1H), 0.93 – 1.09 (m, 1H), 0.40 – 0.54 (m, 2H), 0.12 – 0.24 (m, 2H); ^13^C NMR (DMSO-*d_6_*) δ 167.37, 160.94, 153.08, 143.09, 136.74, 129.33, 126.46, 117.17, 113.84, 107.79, 103.06, 75.18, 71.71, 71.16, 68.40, 68.16, 67.69, 65.26, 52.37, 48.24, 10.38, 2.86, 2.84; RP-HPLC *t*_R_: 2.03 min (System 1), 2.17 min (System 5). *m/z* MS (TOF ES^+^) C_25_H_33_N_2_O_4_ [MH]^+^ calcd 473.2; found 473.3; *m/z* HRMS (TOF ES^+^) C_25_H_33_N_2_O_7_ [MH]^+^ calcd 473.2282; found 473.2215.

*5-(2-((S)-3-((R)-2-((Cyclopropylmethoxy)methyl)-2,3-dihydrobenzo[b][1,4]dioxin-6-yloxy)-2-hydroxypropylamino)ethoxy)isoindolin-1-one (**NDD825**).* Prepared by general method A using 5-(2-Aminoethoxy)isoindolin-1-one hydrochloride (**11**). Yield: 30%, off-white solid; Mp = 159-162 °C; ^1^H NMR (DMSO-*d*_6_) δ 8.29 (s, 1H), 7.55 (d, *J* = 8.4 Hz, 1H), 7.11 (d, *J* = 2.2 Hz, 1H), 7.01 (dd, *J* = 8.4, 2.2 Hz, 1H), 6.77 (d, *J* = 8.8 Hz, 1H), 6.45 (d, *J* = 2.9 Hz, 1H), 6.41 (dd, *J* = 8.8, 2.9 Hz, 1H), 5.00 (s, 1H), 4.30 (d, *J* = 1.7 Hz, 2H), 4.27 (d, *J* = 2.4 Hz, 1H), 4.23 (dtd, *J* = 7.5, 5.1, 2.4 Hz, 1H), 4.10 (t, *J* = 5.5 Hz, 2H), 3.96 (dd, *J* = 11.3, 7.4 Hz, 1H), 3.89 – 3.73 (m, 3H), 3.62 (h, *J* = 5.5 Hz, 2H), 3.29 (d, *J* = 6.8 Hz, 2H), 2.93 (t, *J* = 5.5 Hz, 2H), 2.73 (dd, *J* = 11.8, 4.1 Hz, 1H), 2.63 (dd, *J* = 11.9, 6.3 Hz, 1H), 1.09 – 0.92 (m, 1H), 0.54 – 0.40 (m, 2H), 0.21 – 0.13 (m, 2H); ^13^C NMR (DMSO-*d*_6_) δ 169.8, 161.5, 153.1, 146.5, 143.1, 136.7, 125.1, 124.0, 117.2, 115.1, 108.6, 107.8, 103.1, 75.2, 71.7, 71.1, 68.4, 68.1, 67.9, 65.3, 52.3, 48.2, 44.7, 40.1, 39.9, 39.7, 39.5, 39.3, 39.1, 38.9, 10.4, 2.9, 2.8; RP-HPLC *t*_R_: 2.01 min (System 1), 2.18 min (System 5). *m/z* MS (TOF ES^+^) C_26_H_33_N_2_O_7_ [MH]^+^ calcd 485.2; found 485.4; *m/z*HRMS (TOF ES^+^) C_26_H_33_N_2_O_7_ [MH]^+^ calcd 485.2282; found 485.2290.

### Cloning, expression and purification of tβ_1_AR

A construct encoding tβ_1_AR followed by a thrombin cleavage site and an His8 tag (tβ_1_AR-thrombin-His8) was cloned into pAcGP67B (BD Biosciences). The construct contained the following mutations: N-terminal deletion (ΔM1–V31), ICL3 deletion (ΔC244–R271), C-terminal deletion (Δ365–483), mutations to increase expression (C116L, E130W), mutations in ECL2 found in hβ_1_AR (D184A, D186S, P187D, Q188E, L190R, K191R), a thermostabilising mutation (M281F) and a mutation to remove the palmitoylation site (C358A). High-titre recombinant baculoviruses were prepared using the flashBAC ULTRA system (Oxford Expression Technologies). *Trichoplusia ni* High Five cells (Thermo Fisher Scientific) were grown in suspension in ESF 921 medium (Expression Systems). Insect cells at 2 × 10^6^ cells/mL were infected with 2.5% (v/v) tβ_1_AR baculovirus and cultured for 48 h.

The cells were harvested by centrifugation, flash-frozen in liquid nitrogen, and stored at −80 °C until further use. The membrane fraction was prepared through two iterations of homogenization using an Ultra-Turrax disperser (IKA) and centrifugation at 40,000g for 1 h in 20 mM Tris pH 7.4, 350 mM NaCl, and 1 mM EDTA. The membranes were solubilised in 1.7% (w/v) lauryl maltose neopentyl glycol (LMNG, Anatrace) and further purified with 0.005% LMNG by Ni^2+^-affinity chromatography as previously described^47^. The eluate was concentrated using a 100 kDa molecular weight cut-off (MWCO) Amicon Ultra centrifugal concentrator (Merck) and further polished on a Superdex 200 Increase column, pre-equilibrated in 20 mM Tris pH 7.4, 500 mM NaCl, 0.005% (w/v) LMNG, and 0.2 mM EDTA. The peak fractions containing β_1_AR were pooled and concentrated.

### Engineering, cloning, expression and purification of Nb60.5

The Nb60.5 variant was identified through Deep Screening^34^. Two focused libraries were designed based on Nb60^33^, incorporating single (Asn32) and double (Leu31 and Asn32) deletions within the CDR1 region, resulting in a total of 6.6 million unique variants that were synthesised by Twist Bioscience. Following a single round of Deep Screening, six nanobody clones were selected for further characterisation, each exhibiting nanomolar binding affinity as confirmed by Octet biolayer interferometry (BLI) analysis. Nb60.5 was chosen for structural studies.

A synthetic gene (Integrated DNA Technologies) for Nb60.5 was cloned into plasmid pET-26b(+) (Novagen) with an N-terminal His6 tag followed by a thrombin protease cleavage site. Expression in E. coli strain BL21(DE3) RIL (Agilent Technologies) and purification from the periplasmic fraction were as described elsewhere^48^. The His6 tag was removed by a thrombin (Sigma) protease cleavage step before concentrating the nanobody to 49 mg/mL.

### Formation of ligand-bound tβ_1_AR-Nb60.5 complex

Preparation of the receptor-nanobody complex was performed as described previously^45^. tβ_1_AR (0.8 mg) was mixed with a 2.5-fold molar excess of nanobody (1 mg), and CGP20712A, NDD713, or NDD825 was added to a concentration of 0.5 mM in a final volume of 100 μL. The mixture was then incubated for 2 h on ice. After incubation, size-exclusion chromatography (SEC) was performed to separate the receptor–nanobody complex from excess nanobody. The complex was loaded and separated on a Superdex 200 10/300 GL column (GE Healthcare), pre-equilibrated with 20 mM Tris-HCl (pH 7.4), 500 mM NaCl, 0.2 mM EDTA, and 0.005% LMNG. The peak fractions corresponding to the complex were concentrated to 12 mg/mL.

### Ligand-bound tβ_1_AR-Nb60.5 complex cryo-grid preparation and data collection

Cryo-EM grids were prepared by applying 3 μL sample (at a protein concentration of 12 mg/mL) on plasma-treated holey gold grids (UltrAufoil Au 1.2/1.3 300 mesh). Excess sample was removed by blotting with filter paper for 4 s before plunge-freezing in liquid ethane (cooled to −181 °C) using a FEI Vitrobot Mark IV maintained at 100% relative humidity and 4 °C. Data collection was carried out on grids made from a single preparation of tβ_1_AR-Nb60.5 complex. Images were collected on a FEI Titan Krios microscope at 300 kV using a Falcon 4i electron detector in counting mode. Data were collected in three independent sessions on LMB Krios2 (CGP20712A and NDD713) and LMB Krios4 (NDD825) using EPU (Thermo Fisher Scientific). Individual grid holes were centred with stage shift for data acquisition. In all sessions, three non-overlapping exposures were collected per grid hole. Micrographs were collected with a fluence of 60 e^−^/Å^2^. Each micrograph was collected as a series of movie frames (approximately 1.0 e^−^/Å^2^/frame) at a flux of 6.7 e^−^/pixel/s (LMB Krios2) or 7.3 e^−^/pixel/s (LMB Krios4), with an energy selection slit width of 20 eV. The datasets were collected at a magnification of 120,000× (0.67 Å/pixel, LMB Krios2) and 165,000× (0.744 Å/pixel, LMB Krios4). The gain reference file was provided by the facility and used unmodified.

### Cryo-EM data processing, model building and refinement

CryoSPARC v4.5.3^49^ was used for all data processing unless otherwise specified. Overall, drift, beam-induced motion and dose weighting were corrected with Patch Motion Corr. Contrast Transfer Function (CTF) fitting and phase shift estimation were performed using Patch CTF estimation. The exposures were manually curated: the only images kept had an estimated CTF resolution of < 5 Å. Autopicking was performed using Blob Picker. Particle picks were curated to remove obvious junk picks by extraction with a box size of 360 pixels and then down-sampled four times. The particles were subjected to two rounds of 2D classification, and the clean particle stack was selected. These sorted particles were applied to Ab-Initio Reconstruction and classified into four classes. Iterative Hetero Refinement was performed with the four classes as an input. Particles were chosen from the good class and subjected to 2D Classification. Particles belonging to the clean 2D classes were re-extracted with a box size of 400 pixels (CGP20712A and NDD825) and processed for Non-Uniform Refinement, yielding high-resolution density maps with a global resolution of 3.0 Å (CGP20712A) and 2.8 Å (NDD825). A subsequent 3D Classification was performed for NDD713, and the particles belonging to the good class were re-extracted with a box size of 320 pixels for Non-Uniform Refinement, reaching a global resolution of 3.3 Å.

The initial tβ_1_AR-Nb60.5 model was generated using ModelAngelo^50^. Ligand CIF files were generated from Acedrg^51^. After docking the models into the corresponding EM density maps using UCSF Chimera^52^, iterative rounds of manual adjustment and real-space refinement were performed in Coot0.9 and real-space refinement in Phenix^53^, respectively. MolProbity^54^ was used to validate the final models, and the comprehensive refining statistics are provided in Extended Data Table 1.

### Determination of ligand affinity for hβ_1_AR and hβ_2_ARand chimeric mutant receptors

The hβ_1_AR and hβ_2_AR constructs have been described previously^22–24^. Receptor DNA sequences containing each of the 5 single point mutations (hβ_1_AR: I118H, V119I, V189T, F359Y or V360I; hβ_2_AR: H93I, I94V, T164V, Y308F or I309V) and receptors with all 5 mutations together (hβ_1_AR•hβ_2_OBS: hβ_1_AR with I118H, V119I, V189T, F359Y and V360I mutations and hβ_2_AR•hβ_1_OBS: hβ_2_AR with H93I, I94V, T164V, Y308F and I309V mutations) in pcDNA3.1 on a neomycin marker were obtained from GeneScript Biotech (UK) Ltd (Oxford, UK). All plasmids used in the study were verified by Oxford Nanopore whole plasmid sequencing (Plasmidsaurus).

CHO-K1 cells (RIDD: CVCL_0214) stably expressing a CRE-SPAP reporter gene^22–24^ were grown in Dulbecco’s modified Eagle’s medium nutrient mix F12 (Sigma Aldrich, Poole, Dorset, UK DMEM/F12: D6421) supplemented with 2 mM L-glutamine (G7513) and 10% (v/v) foetal calf serum (F7524) in a T25 flask in a 37°C humidified 5% CO_2_ : 95% air atmosphere. Cells were transfected with hβ_1_AR, hβ_2_AR or one of the mutant receptors using 6ng plasmid DNA in 30 µl Lipofectamine (Life Technologies; Paisley, UK) and 2.5ml OPTIMEM. After 24 h, the transfection reagents removed and replaced with DMEM/F12, containing 10% (v/v) foetal calf serum, 2mM L-glutamine, neomycin (1 mg/ml) and hygromycin (200 µg/ml). Cells were passaged in this selection media until two confluent T175 flasks of cells were obtained, after which antibiotics were withdrawn. Cells were then plated into tissue-culture-treated white-sided 96-well view plates and grown to confluence. Three separate transfections, yielding 3 separate stable mixed populations of cells for each receptor were used, and 2-3 separate experiments determined from each population giving 6-7 overall separate experiments.

^3^H-CGP12177 whole cell binding in living cells was used to determine the affinity of ligands. The affinity (K_D,_ concentration required to bind half of the receptors) of ^3^H-CGP12177 was determined for each construct from saturation experiments. Cells were incubated for 2 h (37°C humidified 5% CO_2_ : 95% air atmosphere) in 200 µl per well of ^3^H-CGP12177 (0.006 to 12.6 nM; Amersham International, Buckinghamshire, UK) in the absence or presence of 10 µM propranolol (P0884, Sigma Aldrich, Poole, Dorset, UK; used to determine non-specific binding), in quadruplicated wells per experiment. Cells were then washed with 2 x 200 ml 4°C phosphate buffered saline and then 100 µl Microscint 20 (PerkinElmer, Shelton, CT, USA) was added to each well and the plates left for several hours in the dark before being counted on a Topcount (2 min per well). Specific binding (SB, equation 1) of ^3^H-CGP12177 at different concentrations of ^3^H-CGP12177 was fitted using the non-linear regression program Prism 7 to the equation:

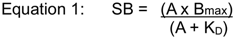

where A is the concentration of ^3^H-CGP12177, B_max_ is the maximal specific binding and thus the K_D_ of ^3^H-CGP12177 was determined.

The IC_50_ value for competing ligands (CGP20712A (Tocris Life Sciences; Avonmouth, UK), NDD713 and NDD825) was determined from a competition curve whereby the competing ligand (over a concentration range of 0.01 nM to 100 µM, 200 µl per well) was incubated (2 h, 37°C) in the presence of a fixed concentration of ^3^H-CGP12177 (range 0.59-0.71 nM) in triplicate wells per experiment. Total and non-specific binding (10 µM propranolol) were determined in 6 replicates in every plate. Cells were washed as above. A sigmoidal concentration-response curve was then fitted to the data using Graphpad Prism 7 and the IC_50_ was determined as the concentration required to inhibit 50% of the specific binding using equation 2.

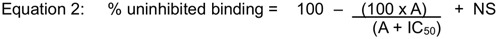

where A in the concentration of the competing ligand, IC_50_ is the concentration at which half of the specific binding of ^3^H-CGP12177 has been inhibited, and NS is the non-specific binding. From the IC_50_ value and the known concentration of ^3^H-CGP12177, a K_D_ value was calculated using equation 3:

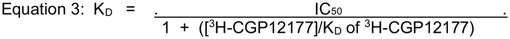

To determine whether a single mutation (or all 5 mutations together) was statistically different from all the others, a one-way ANOVA with post hoc Tukey analysis was conducted. The one-way ANOVA determines if there is a difference within the datasets as a whole, when comparing each mutant with each other and wildtype in a single statistical test. If the ANOVA does detect a difference, the post hoc analysis determines within the entire dataset which mutants have caused a change that is statistically different from wildtype, with a *P* value set at p<0.0001.

### Ligand binding kinetics assay

The hβ_1_AR, hβ_2_AR, hβ_1_AR_out_/hβ_2_AR_in_ and the chimeras hβ_2_AR_out_/hβ_1_AR_in_ were synthesised and cloned into a lentiviral plasmid pHR-CMV-TetO2. The constructs were stably expressed in HEK293 GnTi-TetR cell lines^55^ (ATCC), which allow for tetracycline-inducible expression. The constructs expression was induced by exposure to 100 ng/mL tetracycline (Sigma) for 18 hours. Harvested cells were counted, snap-frozen in liquid nitrogen and then stored at −80°C. For the radioligand assay, generally, cells were thawed and homogenised in lysis buffer (20 mM HEPES-NaOH pH 7.5, 50 mM NaCl, 2.5 mM MgCl_2_, 1 mg/mL BSA). Appropriate dilutions were prepared, and 110 μL of aliquots were loaded into a Multiscreen GF/B filter plate (Millipore) blocked with lysis buffer plus 0.1% PEI. Appropriate ligands were added to each well, mixed and incubated for the appropriate time. The cells were filtered through the plate on a vacuum unit, and the plate was dried at 60°C for 60 min. Filter papers containing the cells were punched into biovials and incubated with 4 mL liquid scintillation cocktail. The radioactivity was counted by a Revvity scintillation counter.

The ligand binding kinetics assay was performed as described previously^42^ with modifications. In general, the [^3^H]DHA dissociation binding kinetics were measured by incubating cells with 5 nM [^3^H]DHA in lysis buffer for 1 h, followed by the addition of 10 μM propranolol for different times. The amount of [^3^H]DHA bound to the filter were measured by scintillation counting, and dissociation rate constants were determined by fitting to a one-phase exponential decay in GraphPad Prism 11. The [^3^H]DHA association binding kinetics were measured by incubating cells with 5 nM [^3^H]DHA for different times, and association rate constants were determined using an association kinetics (one concentration of hot) fit in GraphPad Prism. Displacement experiments were then performed using 8 different concentrations of competing ligands in the presence of 5 nM [^3^H]DHA to determine the inhibitory constant (K_i_) of unlabelled CGP20712A, NDD713, and NDD825 ligands. To determine the binding kinetics of unlabelled ligands, cells were incubated with four different concentrations (0 x *K*i, 0.3 x *K*i, 1 × *K*i, 3 x *K*i) of unlabelled ligands and 5 nM [^3^H]DHA for different times. Binding kinetics were determined using the kinetics of competitive binding fit in GraphPad Prism with the previously measured kinetic parameters for [3H]DHA. Each condition was measured in three independent experiments, each performed in duplicate. Outliers were identified separately for k_on_, k_off_ and *K*_D_ using the ROUT method in GraphPad Prism (Q = 1%), so the number of replicates contributing to each parameter can differ; n is given with each value in Extended Data Table 4.

### Molecular dynamics simulations

Inactive hβ_1_AR and hβ_2_AR models were prepared from PDB entries 7BVQ and 2RH1, respectively, using SWISS-MODEL^56^. Where used, the protonation state of CGP20712A was checked with SwissParam^57^, after which the ligand was processed through the CHARMM-GUI^58^, with the central secondary aliphatic amine modelled in its protonated state to give a net charge of +1. Simulation systems were constructed in CHARMM-GUI using the CHARMM36m^59^ and CGenFF ligand force fields^60^. Each receptor was embedded in a POPC membrane containing 20 mol% cholesterol, solvated with TIP3P water and 0.15 M NaCl. Following energy minimisation, the systems were equilibrated for 1.875 ns using a six-stage CHARMM-GUI membrane protocol with progressively reduced restraints. Unbiased molecular dynamics simulations were performed with GROMACS 2024.4^61^. A 2-fs time step was used, with temperature maintained at 303.15 K using the velocity-rescaling thermostat^62^ and pressure maintained at 1 bar using the semi-isotropic Parrinello–Rahman barostat^63^. Five replicate 500 ns apo simulations were performed for each receptor.

For umbrella sampling, potentials of mean force (PMFs) were calculated for the separation between H4 and H5 in apo hβ_1_AR and hβ_2_AR. The reaction coordinate, q, was defined as the all-atom centre-of-mass distance between Leu188-Ser190 and Val231-Ser232 in hβ_1_AR, and between Leu163–Ser165 and Val206-Ser207 in β_2_AR. Sixteen windows were distributed uniformly between q = 0.80 - 1.12 nm and restrained with a harmonic force constant of 1,000 kJ/(mol·nm²)^64^. Starting conformations were selected directly from the unbiased apo simulations without a pulling simulation. Each window was simulated for 20 ns, with the first 2 ns excluded from downstream analysis. PMFs were reconstructed over 2-20 ns using gmx wham in GROMACS 2023.4 at 303.15 K^65^. Each profile was referenced to its global minimum. Uncertainty was estimated from 200 histogram-bootstrap profiles. Histogram overlap and PMFs reconstructed over 2–10, 2–15 and 2–20 ns were used to assess sampling and time dependence (Extended Data Fig. 11a-b).

Multiple-walker well-tempered metadynamics (mwWT-MetaD) simulations were performed using GROMACS 2022.4 patched with PLUMED 2.8.2^66–68^. Simulations were initiated from the inactive receptors with a closed H4-H5 region, with CGP20712A placed in the extracellular solvent to avoid biasing the initial opening of this region. Sixteen walkers from the same input pose were used per receptor, providing aggregate sampling times of 14.83 μs for hβ_1_AR and 14.27 μs for β_2_AR.

The biased collective variable, d, was the centre-of-mass distance between the three fluorine atoms of the ligand and two receptor Cα atoms: Val189 and Val231 in hβ_1_AR, and Thr164 and Val206 in β_2_AR. Gaussian potentials with an initial height of 1.0 kJ/mol and a width of 0.05 nm were deposited every 1 ps using a bias factor of 20 at 303.15 K. One-dimensional free-energy profiles were reconstructed using plumed sum_hills with 600 bins and referenced to the median free energy of the fully solvated region. The resulting profiles were interpreted as relative free energies along d. Convergence was assessed from time-resolved profiles and Gaussian-height decay (Extended Data Fig. 11c-d). For each free-energy minimum, frames within 0.1 nm of the minimum were clustered based on RMSD after protein alignment, and the centroid of the most populated ligand heavy-atom cluster was selected as the representative structure.

## Data Availability Statement

The cryo-EM density maps have been deposited in the Electron Microscopy Data Bank (EMDB) under accession codes EMD-100246, EMD-100249 and EMD-100274 (CGP20712A, NDD713 and NDD825 bound to tβ_1_AR-Nb60.5, respectively). The atomic coordinates have been deposited in the Protein Data Bank (PDB) under accession codes PDB 34YB, 34YG and 34YR (CGP20712A, NDD713 and NDD825 bound to tβ_1_AR-Nb60.5, respectively). All input tpr files along with secondary analysis and selected snapshots from the MD data have been deposited in OSF and are freely available at https://doi.org/10.17605/OSF.IO/TEV5C. There are no restrictions on data availability and they are available on reasonable request.

## Acknowledgements

We acknowledge the MRC Laboratory of Molecular Biology Electron Microscopy Facility for access and support of electron microscopy sample preparation and data collection, the Baculovirus and Mammalian TC facility for insect cell preparation, the LMB scientific computing for technical support and the Flow Cytometry facility for support in cell analysis. The work in C.G.T.’s laboratory was supported by core funding from the Medical Research Council [MC_U105197215]. The work was also supported by the Medical Research Council (MRC) program grant MC_U105178804 (P.H.), a research collaboration between AstraZeneca UK Limited and the Medical Research Council – MRC-AstraZeneca Blue Sky Grant (B.T.P.) and a Wellcome Trust Seeding Drug Discovery Initiative Grant 086039/Z/08/Z (JGB). RAC is supported by a MRC project grant (APP54028). XX is supported by a China Scholarship Council (CSC) PhD. This project made use of time on ARCHER2 granted via the UK High-End Computing Consortium for Biomolecular Simulation, HECBioSim (http://www.hecbiosim.ac.uk), supported by EPSRC (grant no. EP/R029407/1). This work was carried out using the computational facilities of the Advanced Computing Research Centre, University of Bristol - http://www.bristol.ac.uk/acrc/. For the purpose of open access, the MRC Laboratory of Molecular Biology has applied a CC BY public copyright licence to any Author Accepted Manuscript version arising.

## Author contributions

Q.C. performed receptor and nanobody, purification, preparation of cryo-EM grids, cryo-EM data collection, data processing, structure determination and model building. Q.C. and C.G.T. carried out structure analysis. Q.C. cloned receptor mutants, made stable cell lines and performed kinetic analyses. T.W. and P.C.E. performed receptor and nanobody purifications in the initial stages of the project. B.T.P. performed the nanobody library design and Deep Screening and P.H. supervised. X.X. performed MD simulations, supervised by R.A.C. and S.J.M., and X.X., R.A.C., and S.J.M. analysed the data. S.N.M., G.P.J. and C.F. synthesised NDD713 and NDD825 and P.M.F and B.K. supervised. J.G.B. designed, performed and analysed the whole cell binding studies with tβ_1_AR, hβ_1_AR and hβ_2_AR and their mutants, and S.J.H. supervised. S.N.M. advised on the stereochemistry of NDD713 and NDD825 and structural modelling. Q.C. and C.G.T. wrote initial drafts of the manuscript and all authors contributed to the manuscript in its final form. C.G.T. managed the overall project.

