## Supplementary figures and tables for "Receptor dynamics and kinetic discrimination contribute to selectivity of β_1_-adrenoceptor antagonists"

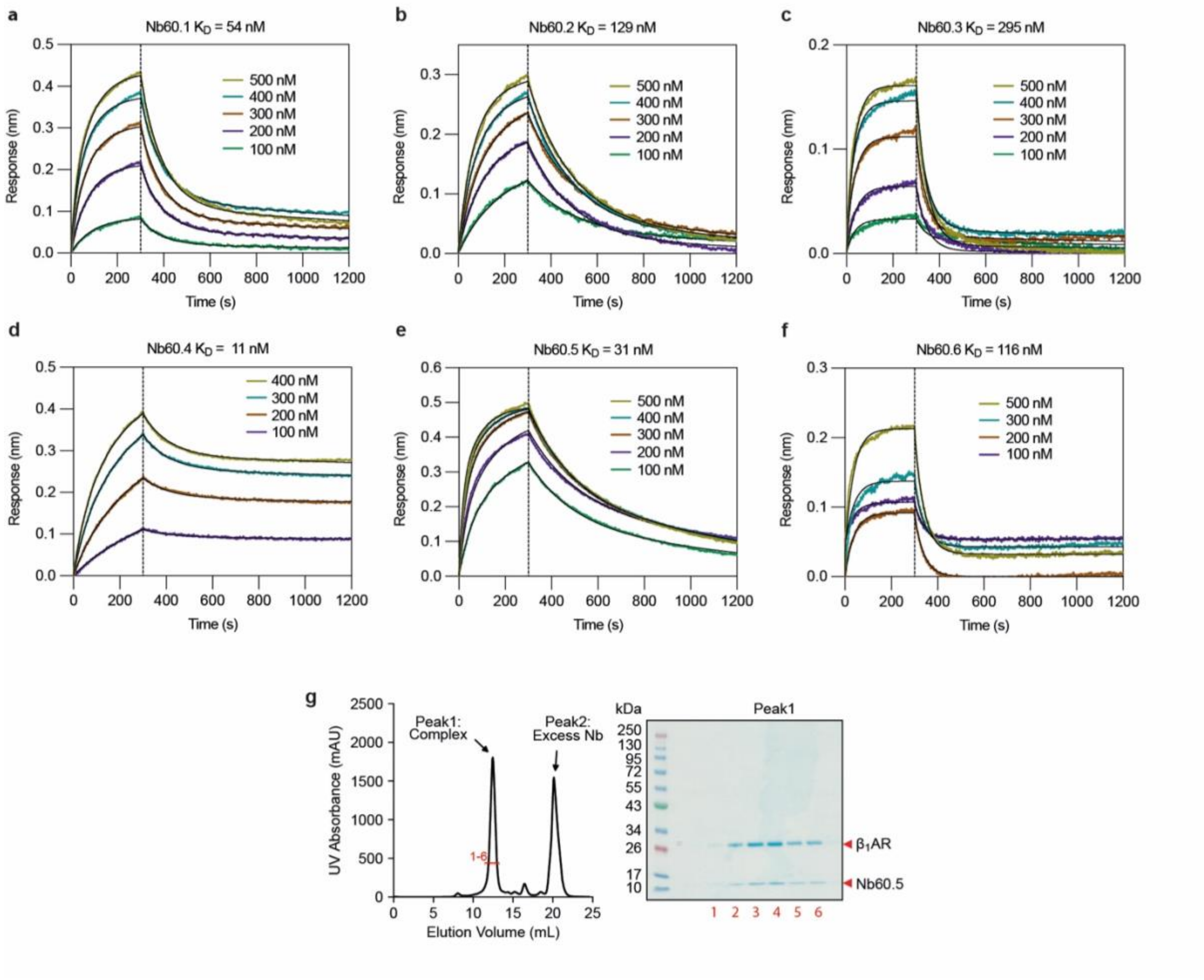

**Extended Data Fig. 1 | Binding kinetics of the selected nanobodies Nb60.1-Nb60.6 to t$\beta_1$AR. a-f**, Biolayer interferometry sensorgrams for Nb60.1 (**a**), Nb60.2 (**b**), Nb60.3 (**c**), Nb60.4 (**d**), Nb60.5 (**e**) and Nb60.6 (**f**) binding to t$\beta_1$AR. Coloured traces are the measured responses at the nanobody concentrations; black lines are global fits to a 1:1 binding model that were used subsequently to determine $K_D$s. The dashed line marks the start of dissociation. Data are from a single experiment. **g**, Size exclusion chromatographic trace (left) of the t$\beta_1$AR-Nb60.5 complex and associated Coomassie Blue-stained SDS-PAGE gel of six fractions across Peak 1.

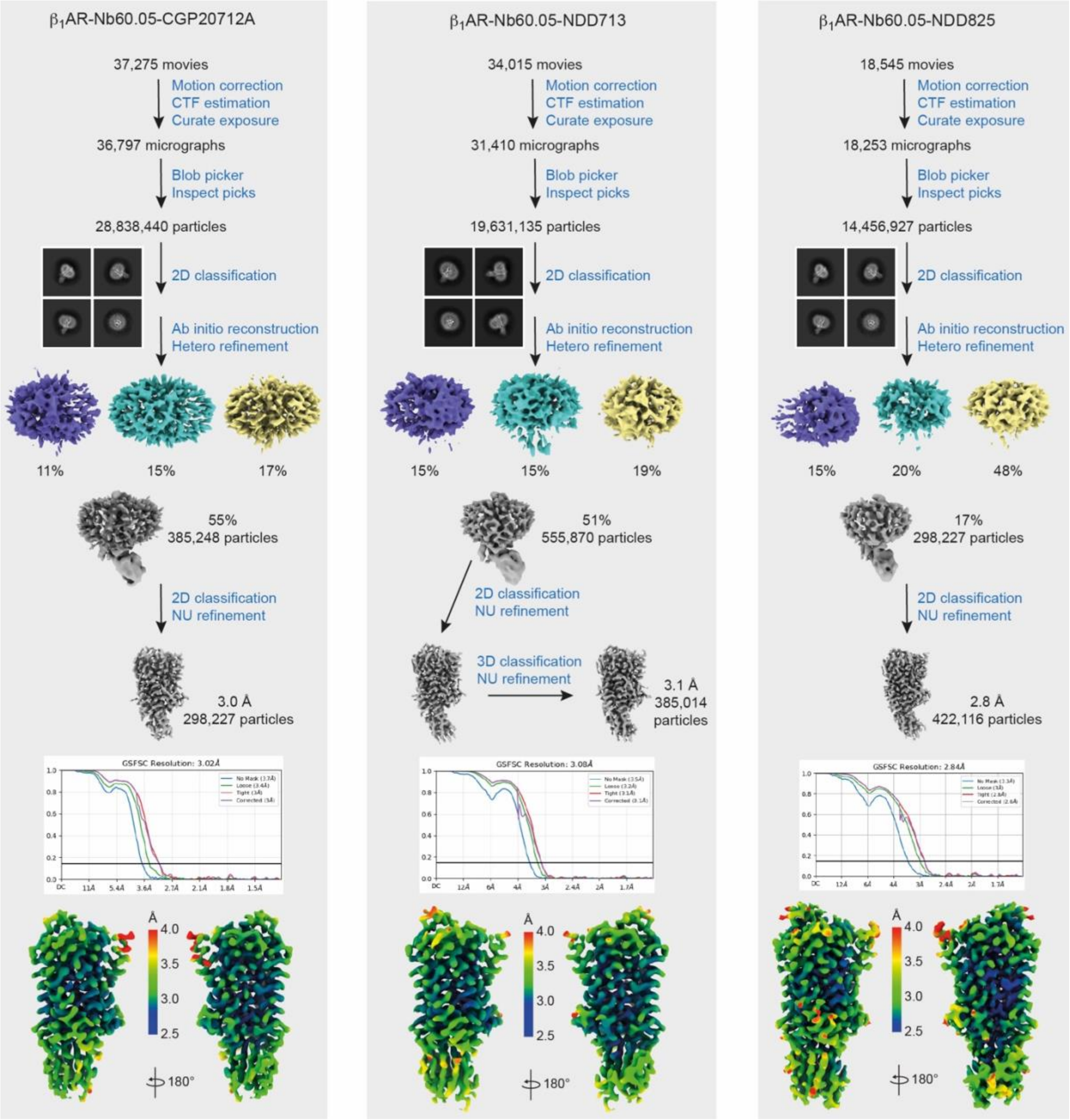

**Extended Data Fig. 2 | Cryo-EM data processing of the t$\beta_1$AR-Nb60.5 complexes bound to CGP20712A, NDD713 and NDD825. a-c**, Cryo-EM data processing workflows for t$\beta_1$AR-Nb60.5 bound to CGP20712A (**a**), NDD713 (**b**) and NDD825 (**c**). The numbers of movies, micrographs and particles are given at each step. Representative 2D class averages are shown. Gold-standard Fourier shell correlation (GSFSC) curves for the final maps are shown along with local resolution maps at the bottom of each panel.

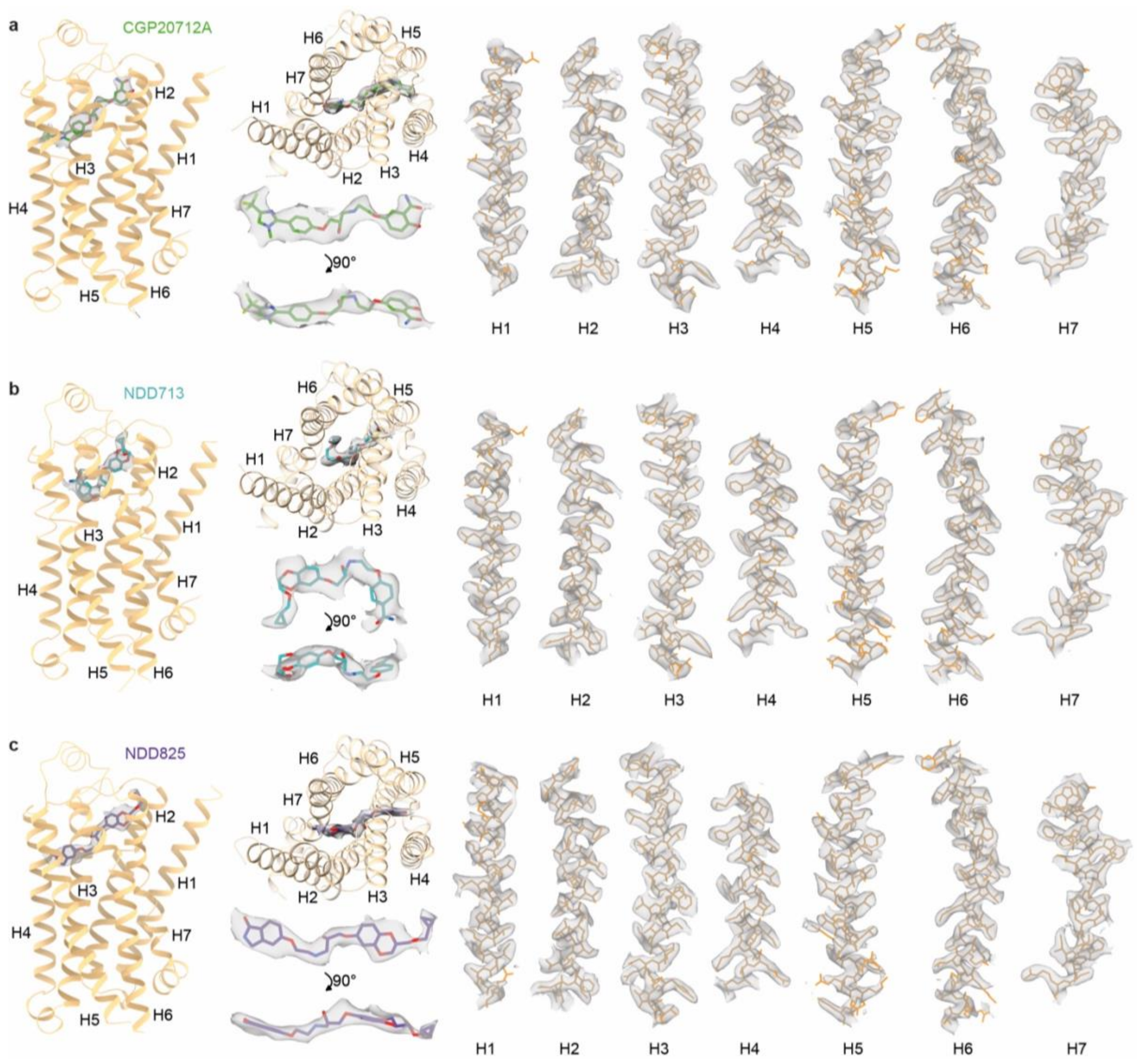

**Extended Data Fig. 3 | Details of density maps and fitted models. a**, CGP20712A-bound t$\beta_1$AR-Nb60.5. **b**, NDD713-bound t$\beta_1$AR-Nb60.5. **c**, NDD825-bound t$\beta_1$AR-Nb60.5. Nanobody Nb60.5 has been omitted for clarity. Panels from left to right show the following: view of t$\beta_1$AR (pale brown) viewed in parallel to the membrane plane, view of t$\beta_1$AR viewed perpendicular to the membrane plane from the extracellular surface, two views of the ligands with associated density, each of the transmembrane $\alpha$-helices.

| Receptor | PDB code | Ligand name | Ligand type | TM2 | | | | EL1 | TM3 | | | | | | TM4 | EL2 | | | | | TM5 | | | | | TM6 | | | | TM7 | | | |
|---|---|---|---|---|---|---|---|---|---|---|---|---|---|---|---|---|---|---|---|---|---|---|---|---|---|---|---|---|---|---|---|---|---|
| | | | | 2.53 | 2.61 | 2.64 | 2.65 | 23.50 | 3.28 | 3.29 | 3.32 | 3.33 | 3.36 | 3.37 | 4.56 | 45.50 | 45.51 | 45.52 | N/A | 5.38 | 5.39 | 5.42 | 5.43 | 5.46 | 6.48 | 6.51 | 6.52 | 6.55 | 7.35 | 7.36 | 7.39 | 7.40 | 7.43 |
| tβ1AR | 2VT4 | (S)-Cyanopindolol | Ant | | | | | | W | T | D | V | V | T | | | | F | T | Y | A | S | S | S | W | F | F | N | | | N | | Y |
| | 2Y01 | Dobutamine | Ag | | | L | V | | W | T | D | V | V | | | | | F | | Y | A | S | S | S | | F | F | N | | V | N | W | Y |
| | 2Y02 | Carmoterol | Ag | | | | | | W | T | D | V | V | | | | | F | T | Y | A | S | S | S | W | F | F | N | F | V | N | W | Y |
| | 2Y03 | (R)-Isoproterenol | Ag | | | | | | W | T | D | V | V | | | | | F | | Y | | S | S | S | W | F | F | N | | | N | | Y |
| | 2Y04 | Levosalbutamol | Ag | | | | | | W | T | D | V | V | T | | | | F | | Y | A | S | S | S | W | F | F | N | | | N | | Y |
| | 2YCW | Carazolol | Ant | | | | | | W | T | D | V | V | T | | | | F | T | Y | A | S | S | S | W | F | F | N | | | N | | Y |
| | 2YCZ | Iodocyanopindolol | Ant | | | | | | W | T | D | V | V | T | | | | F | T | Y | A | S | S | S | W | F | F | N | | | N | | Y |
| | 3ZPQ | CHEMBL200234 | Frag | | | | | | | | D | V | V | T | | | | F | | Y | | S | S | S | W | F | F | N | | | N | | Y |
| | 3ZPR | CHEMBL1559535 | Frag | | | | | | | | D | V | V | | | | | F | T | Y | A | S | S | S | W | F | F | N | | | N | | Y |
| | 4AMI | Bucindolol | Ant | | | L | | | W | T | D | V | V | T | | C | D | F | | Y | | S | S | S | W | F | F | N | F | V | N | | Y |
| | 4AMJ | (S)-Carvedilol | Ant | | G | L | V | | W | | D | V | V | T | | | | F | T | Y | A | S | S | S | W | F | F | N | F | V | N | W | Y |
| | 5A8E | 7-methylcyanopindolol | Ant | | | | | | W | T | D | V | V | T | | | | F | T | Y | A | S | S | S | W | F | F | N | F | | N | | Y |
| | **34YB** | **CGP20712A** | Ant | | | L | V | | W | | D | V | | | V | C | D | | | Y | | S | | S | | F | F | | | | N | W | Y |
| | **34YG** | **NDD713** | Ant | | | L | | | W | | D | V | | | | C | D | F | | Y | | S | | | | F | F | N | F | V | N | | Y |
| | **34YR** | **NDD825** | Ant | | | L | V | | W | T | D | V | | T | | | | | | | | S | | S | | F | | | | V | N | W | Y |
| hβ1AR | 7BVQ | Carazolol | Ant | | | | | | W | T | D | V | V | T | | | | F | T | Y | A | S | S | S | W | F | F | N | | | N | | Y |
| hβ2AR | 3D4S | Timolol | Ant | M | | | | | W | T | D | V | V | T | | | | F | | Y | A | S | S | S | W | F | F | N | Y | | N | | Y |
| | 3NY8 | CHEMBL1233766 | Ant | M | | | | | W | T | D | V | V | T | | | | F | | Y | | S | S | S | W | F | F | N | Y | | N | | Y |
| | 3NY9 | CHEMBL1233771 | Ant | M | | | | | W | T | D | V | V | T | | | | F | T | Y | A | S | S | S | W | F | F | N | Y | | N | | Y |
| | 3NYA | CHEMBL1160734 | Ant | M | | | | | W | T | D | V | V | T | | | | F | | Y | A | S | S | S | W | F | F | N | Y | | N | | Y |
| | 3PDS | FAUC50 | Ag | | G | C | I | W | W | T | D | V | V | | | C | D | F | T | Y | A | S | S | S | W | F | F | N | Y | I | N | W | Y |
| | 5D5A | Carazolol | Ant | | | | | | W | T | D | V | V | T | | | | F | T | Y | A | S | S | S | W | F | F | N | Y | | N | | Y |
| | 6PS3 | (S)-Carvedilol | Ant | | G | C | I | | W | T | D | V | V | T | | | D | F | T | Y | A | S | S | S | W | F | F | N | Y | I | N | W | Y |
| | 6PS5 | Propranolol | Ant | | | | | | W | T | D | V | V | T | | | | F | | Y | A | S | S | S | W | F | F | N | Y | | N | | Y |

**Extended Data Fig. 4 | Amino acid residues making contacts to ligands in inactive state structures of tβ1AR, hβ1AR and hβ2AR.** Ligands are classified as either agonists (Ag) or antagonists (Ant) with residues making ligand contacts (distance ≤ 3.9 Å) shown as coloured squares with the residue given in single letter code. Colours refer to the characteristics of the residue: red, acidic; green aromatic, yellow hydrophobic, purple, hydrophilic; olive green, Cys (intermediate properties).

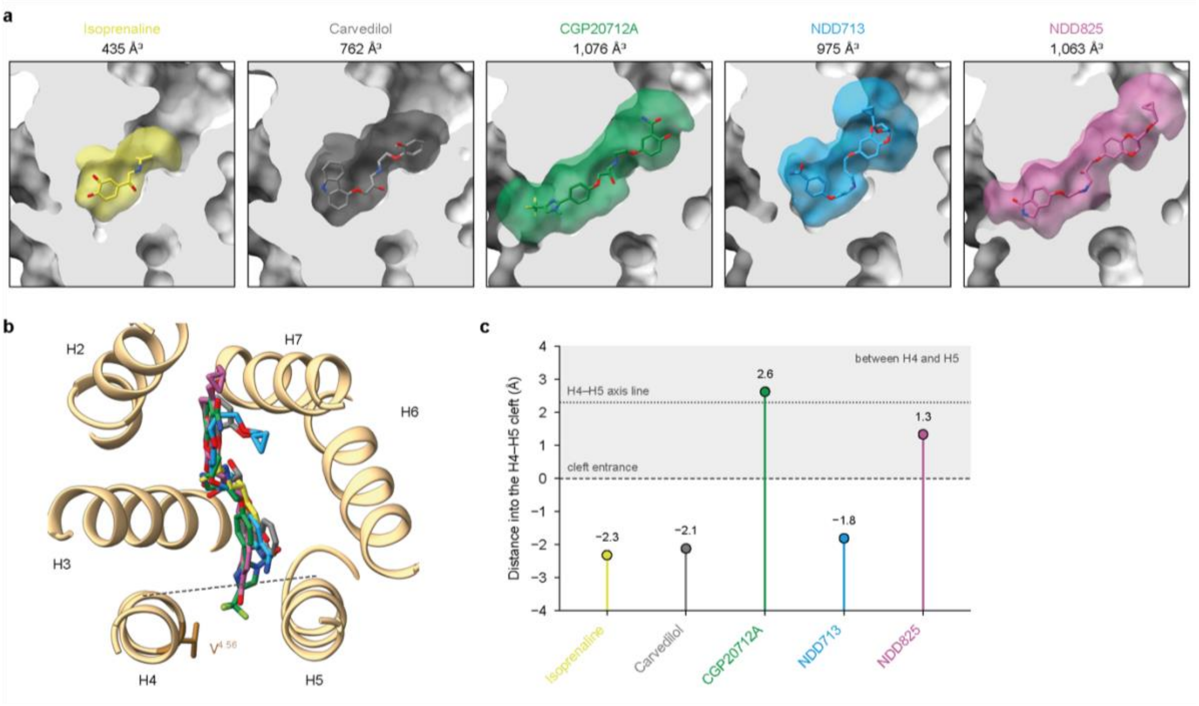

**Extended Data Fig. 5 | Orthosteric binding site occupancy by t$\beta_1$AR ligands and their extension into the H4-H5 cleft. a**, The orthosteric pocket of t$\beta_1$AR bound to isoprenaline (PDB 2Y03), carvedilol (PDB 4AMJ), CGP20712A, NDD713 and NDD825. Each receptor is shown as a molecular surface (grey), clipped through the ligand to expose the OBS; ligands are shown as sticks (carbon in yellow, grey, green, blue and magenta, respectively; nitrogen, blue; oxygen, red; fluorine, light green). The coloured surface marks the space within 4 Å of the ligand (dark grey for carvedilol); its volume is given below each ligand name. All structures were superposed on chain A of the CGP20712A-bound structure and are shown in the same orientation, viewed from the membrane plane with the extracellular side at the top. **b,** Superposition of the five ligands in the CGP20712A-bound t$\beta_1$AR structure, viewed from the extracellular side. The dashed line marks the entrance of the cleft between H4 and H5, defined at the height of the CGP20712A trifluoromethyl group. **c,** Depth to which each ligand reaches into the H4-H5 cleft, measured for its deepest atom. The dashed line (0 Å) marks the cleft entrance, and the dotted line is the line joining the H4 and H5 helix axes; positive values (shaded) lie between H4 and H5. Depth was measured in the membrane plane, using the H4 (residues 165–178) and H5 (residues 205-220) axes of each structure, taken at the height of each ligand atom. The cleft entrance was placed 2.3 Å (the Cα radius of an α-helix) from the axis line, towards the pocket.

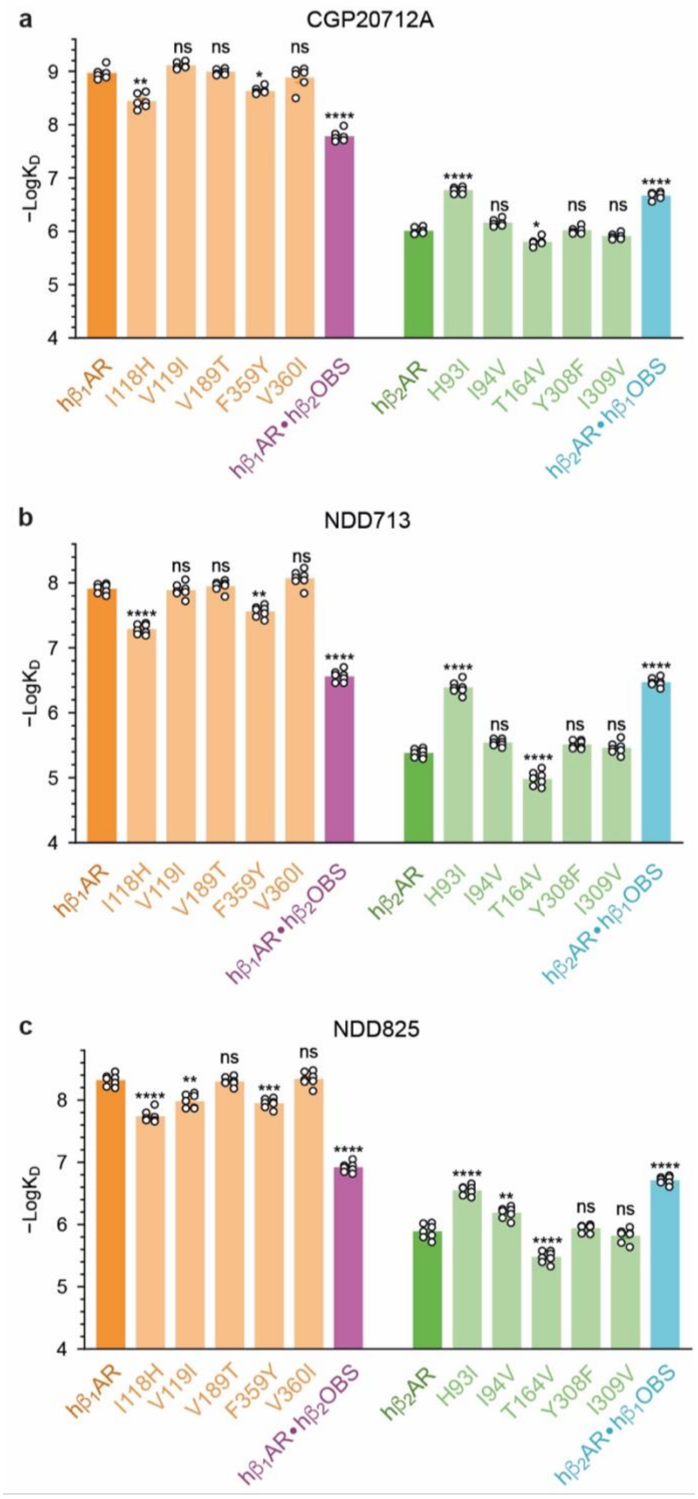

**Extended Data Fig. 6 | Affinities of hβ$_1$AR and hβ$_1$AR mutants**. Affinities (apparent $K_D$ values) for CGP20712A (**a**), NDD713 (**b**) and NDD825 (**c**), binding to hβ$_1$AR (orange), hβ$_1$AR single point mutants (pale orange), hβ$_2$AR (green), hβ$_2$AR single point mutants (pale green), hβ$_1$AR•hβ$_2$OBS (purple) and hβ$_2$AR•hβ$_1$OBS (blue). Affinities are the mean ± sem of 6-7 separate experiments, each performed in triplicate; see Extended Data Table 3 for data and *P* values for comparisons with either hβ$_1$AR or hβ$_2$AR (****, $P \leq 10^{-9}$; ***, $P \leq 10^{-7}$; **, $P \leq 10^{-5}$; *, $P \leq 10^{-3}$; ns, $P > 10^{-3}$).

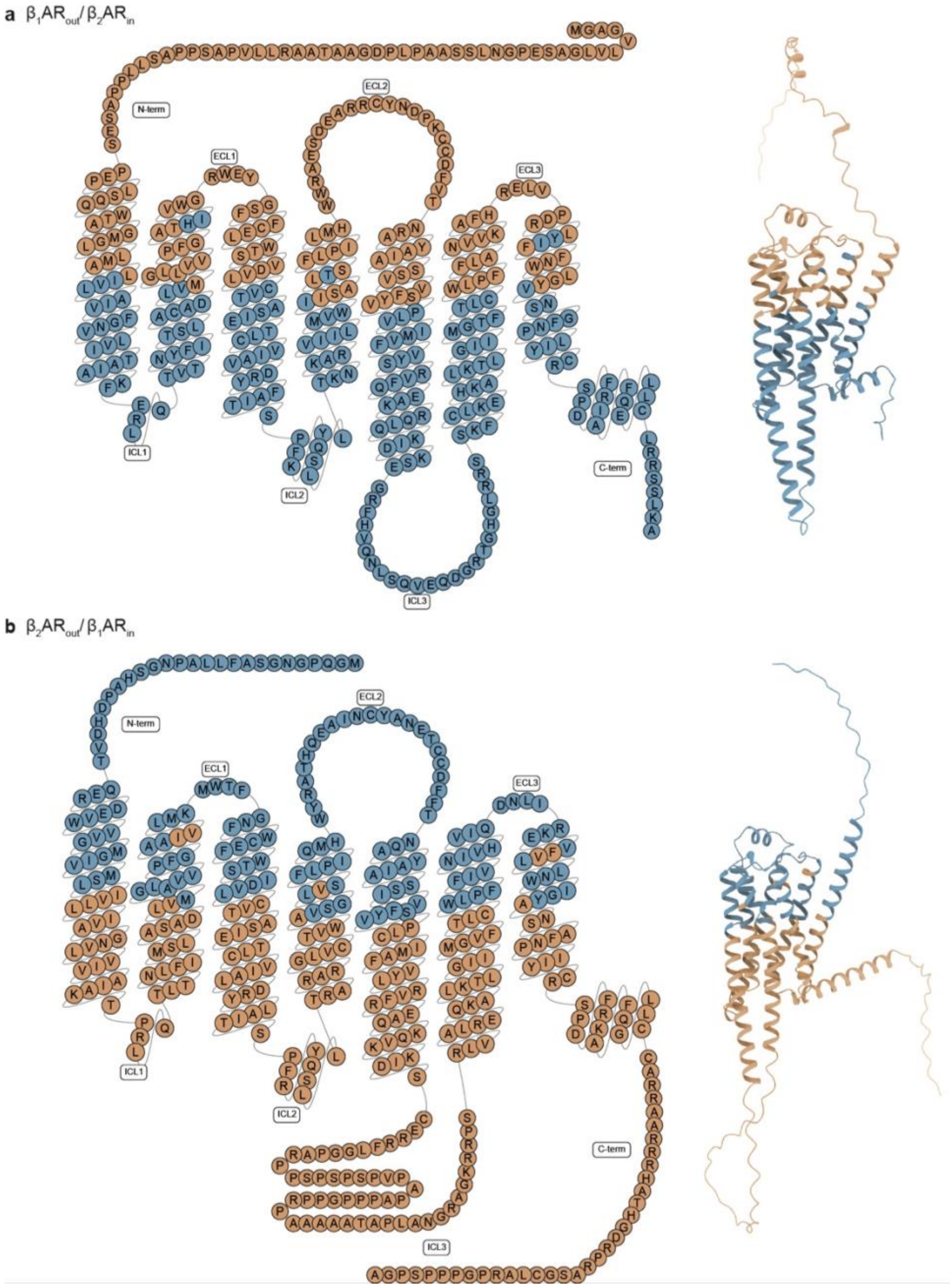

**Extended Data Fig. 7 | Design of the h$\beta_1$AR-h$\beta_2$AR chimeric receptors. a-b,** Snake plot (left) and AlphaFold model (right) of the chimeras h$\beta_1$AR$_{out}$/h$\beta_2$AR$_{in}$ (**a**) and h$\beta_2$AR$_{out}$/h$\beta_1$AR$_{in}$ (**b**). Amino acids are shown in single letter nomenclature in h$\beta_1$AR (brown) and h$\beta_2$AR (blue). The AlphaFold models on the right are in the same colour scheme.

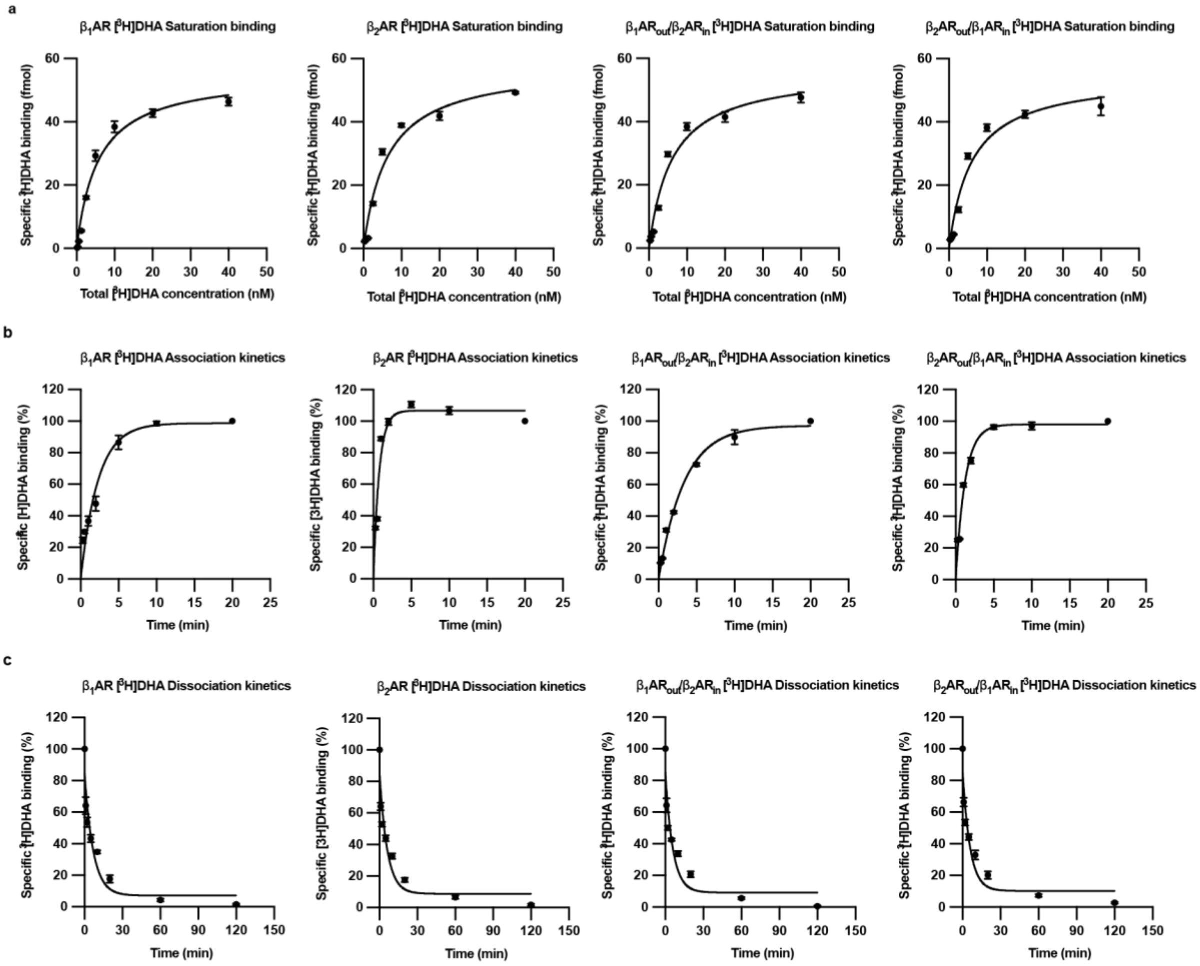

**Extended Data Fig. 8 | [$^3$H]DHA binding to hβ$_1$AR, hβ$_2$AR and the hβ$_1$AR-hβ$_2$AR chimeric receptors. a**, Saturation binding of [$^3$H]DHA. Specific binding is plotted against total [$^3$H]DHA concentration and fitted to a one-site binding model. **b**, Association of [$^3$H]DHA fitted to a one-phase association model. **c**, Dissociation of [$^3$H]DHA fitted to a one-phase decay model. In each panel, the receptors, from left to right, are hβ$_1$AR, hβ$_2$AR, hβ$_1$AR$_{out}$/hβ$_2$AR$_{in}$ and hβ$_2$AR$_{out}$/hβ1AR$_{in}$, as indicated above each plot. Data are the mean ± SEM of 3-6 replicates from three independent experiments each performed in duplicate.

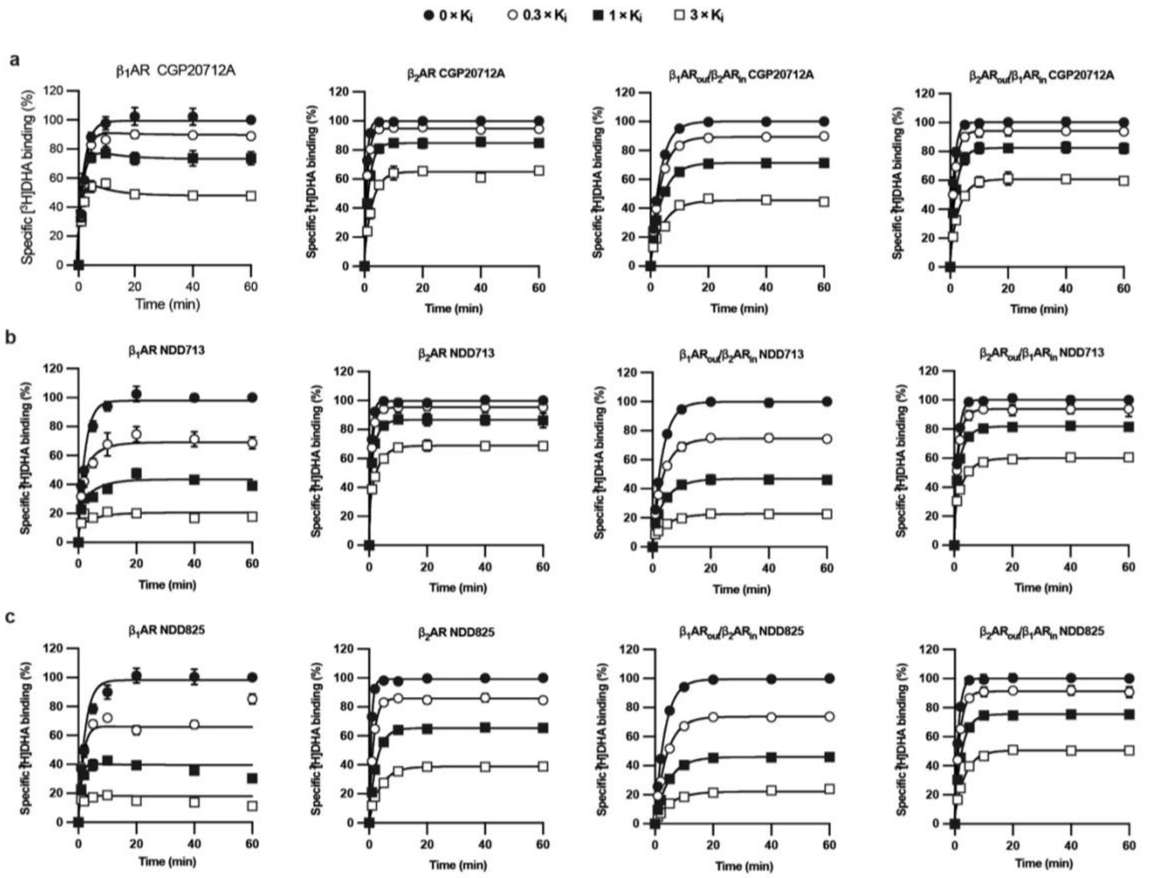

**Extended Data Fig. 9 | Competition association kinetics of CGP20712A, NDD713 and NDD825 at h$\beta_1$AR, h$\beta_2$AR and the h$\beta_1$AR-h$\beta_2$AR chimeric receptors. a-c**, Association of [$^3$H]DHA with h$\beta_1$AR, h$\beta_2$AR, h$\beta_1$AR$_{out}$/h$\beta_2$AR$_{in}$ and h$\beta_2$AR$_{out}$/h$\beta_1$AR$_{in}$ (left to right, as given above each plot) in the absence and presence of CGP20712A (**a**), NDD713 (**b**) and NDD825 (**c**). Solid lines are global fits of all curves in a plot to the Motulsky-Mahan model for competitive binding kinetics. Data are the mean ± SEM of 3-6 replicates from three independent experiments each performed in duplicate. The $k_{on}$, $k_{off}$ and $K_D$ values returned by these fits are those plotted in Fig. 4 and listed in Extended Data Table 4.

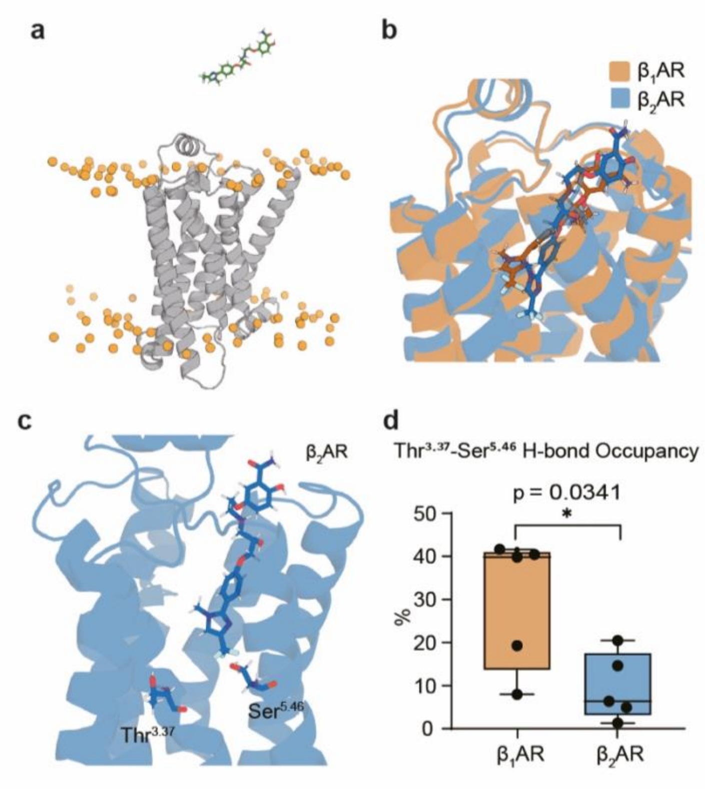

**Extended Data Fig. 10 | Starting conditions for the metadynamics simulations and the Thr$^{3.37}$-Ser$^{5.46}$ hydrogen bond. a,** Starting conformation for the multiple-walker well-tempered metadynamics simulations. The receptor is shown as a grey cartoon and the phosphate atoms of the membrane lipids as orange spheres. **b,** Representative structures of CGP20712A bound to h$\beta_1$AR (light brown) and h$\beta_2$AR (blue) at d = 0.88 nm, superposed on receptor Cα atoms; CGP20712A is shown as sticks in the colour of its receptor. **c,** The interhelical Thr$^{3.37}$-Ser$^{5.46}$ hydrogen bond in h$\beta_2$AR (blue cartoon, residues as sticks). **d,** Occupancy of the Thr$^{3.37}$-Ser$^{5.46}$ hydrogen bond in h$\beta_1$AR (light brown) and h$\beta_2$AR (blue) in five independent apo simulations per receptor, using a distance cutoff of 3.5 Å and an angle cutoff of 30°. Points represent independent simulations. The P-value was calculated from a two-tailed unpaired Student's t-test; *$P < 0.05$.

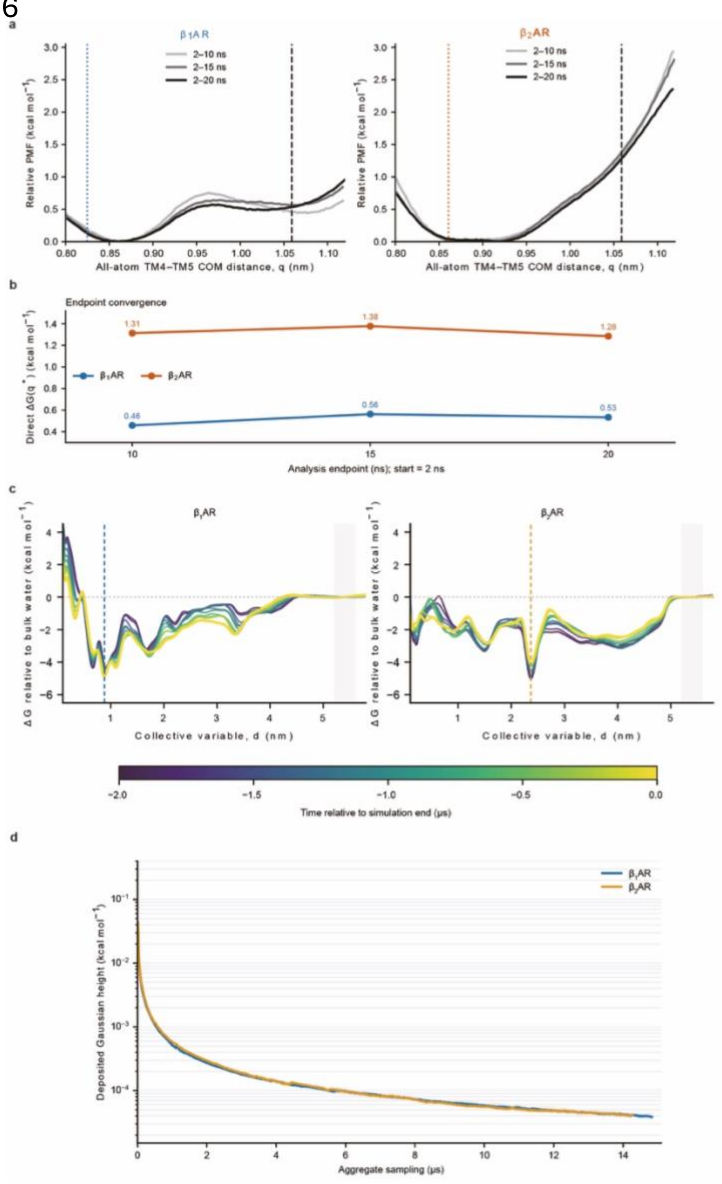

**Extended Data Fig. 11 | Convergence of the umbrella-sampling and metadynamics simulations. a,** Potentials of mean force for H4-H5 separation in hβ$_1$AR (left) and hβ$_2$AR (right), calculated from three analysis windows that all start at 2 ns and end at 10, 15 and 20 ns (greys to black, as in the key). Dotted vertical lines mark q in the corresponding inactive-state structure and the dashed line q* = 1.06 nm, as in Fig. 5b. **b**, Free energy at q* relative to the global minimum, ΔG(q*), for each analysis endpoint, for hβ$_1$AR (pale brown) and hβ$_2$AR (blue); values are given above each point. **c,** Free energy relative to bulk water along the collective variable d for hβ$_1$AR (left) and hβ$_2$AR (right), recalculated at intervals over the last 2 μs of aggregate sampling and coloured by time relative to the end of the simulation (colour bar). Dashed vertical lines mark the global minimum of each profile (0.88 nm for hβ$_1$AR, 2.37 nm for β$_2$AR) and the grey band marks the bulk-water reference region used to set the zero. **d,** Height of the deposited Gaussians against aggregate sampling time for hβ$_1$AR (pale brown) and hβ$_2$AR (blue).

**Extended Data Table 1 | Cryo-EM data collection, refinement and validation statistics**

| | | | |
|---|---|---|---|
| **Data collection and processing** | | | |
| | CGP20712A | NDD713 | NDD825 |
| Magnification | 120,000x | 120,000x | 165,000x |
| Voltage (kV) | 300 | 300 | 300 |
| Electron exposure ($e^-$/Å$^2$) | 60 | 60 | 60 |
| Defocus range (μm) | -0.8 to -1.8 | -0.8 to -1.8 | -0.8 to -1.8 |
| Pixel size (Å) | 0.67 | 0.67 | 0.744 |
| Symmetry imposed | - | - | - |
| Map resolution (Å) | 3.0 | 3.3 | 2.8 |
| FSC threshold | 0.143 | 0.143 | 0.143 |
| **Refinement** | | | |
| Model composition | | | |
| Non-hydrogen atoms | 2833 | 3002 | 2919 |
| Protein residues | 349 | 371 | 360 |
| R.m.s. deviations | | | |
| Bond lengths (Å) | 0.002 | 0.003 | 0.002 |
| Bond angles (°) | 0.552 | 0.520 | 0.512 |
| Validation | | | |
| MolProbity score | 1.17 | 1.29 | 1.28 |
| Clash score | 3.36 | 5.32 | 5.32 |
| Poor rotamers (%) | 0 | 0 | 0 |
| Ramachandran plot | | | |
| Favoured (%) | 97.91% | 98.05 | 98.04 |
| Allowed (%) | 2.09 | 1.95 | 1.96 |
| Disallowed (%) | 0 | 0 | 0 |

### Extended Data Table 2 | Binding affinities (log $K_D$) of CGP20712A, NDD713 and NDD825 at wild-type and chimeric β-adrenoceptors measured in whole cell binding studies

#### a, Data

| Ligand | $t\beta_1AR$ | $h\beta_1AR$ | $h\beta_2AR$ | $h\beta_1AR{\bullet}h\beta_2OBS$ | $h\beta_2AR{\bullet}h\beta_1OBS$ | *n* |
|---|---|---|---|---|---|---|
| CGP20712A | −6.66 ± 0.02 | −8.97 ± 0.05 | −6.01 ± 0.02 | −7.78 ± 0.04 | −6.67 ± 0.03 | 6 |
| NDD713 | −5.25 ± 0.04 | −7.91 ± 0.03 | −5.38 ± 0.02 | −6.56 ± 0.03 | −6.47 ± 0.02 | 7 |
| NDD825 | −5.69 ± 0.04 | −8.32 ± 0.04 | −5.89 ± 0.04 | −6.92 ± 0.03 | −6.71 ± 0.02 | 7 |

Values are log $K_D$ (M), mean ± SEM of *n* independent experiments.

The $K_D$ value for $^3$H-CGP12177 obtained from saturation whole cell binding with $t\beta_1AR$ was 0.27 ± 0.02nM (n=6).

The $K_D$ values for $h\beta_1AR$ (0.28nM) and hβ2AR (0.18nM) were values previous determined in stable mixed populations of cells[22]

#### b, *P* values

| Ligand | $h\beta_1AR$ vs $h\beta_2AR$ | $h\beta_1AR$ vs $h\beta_1AR{\bullet}h\beta_2OBS$ | $h\beta_1AR$ vs $h\beta_2AR{\bullet}h\beta_1OBS$ | $h\beta_2AR$ vs $h\beta_1AR{\bullet}h\beta_2OBS$ | $h\beta_2AR$ vs $h\beta_2AR{\bullet}h\beta_1OBS$ |
|---|---|---|---|---|---|
| CGP20712A | $3 \times 10^{-14}$ | $3 \times 10^{-14}$ | $3 \times 10^{-14}$ | $3 \times 10^{-14}$ | $4 \times 10^{-12}$ |
| NDD713 | $5 \times 10^{-14}$ | $5 \times 10^{-14}$ | $5 \times 10^{-14}$ | $5 \times 10^{-14}$ | $5 \times 10^{-14}$ |
| NDD825 | $5 \times 10^{-14}$ | $5 \times 10^{-14}$ | $5 \times 10^{-14}$ | $5 \times 10^{-14}$ | $05 \times 10^{-14}$ |

*P* values from one-way ANOVA with post hoc Tukey analysis (see methods)

**Extended Data Table 3 | Binding affinities (log $K_D$) of CGP20712A, NDD713 and NDD825 at binding-site mutants of hβ$_1$AR and hβ$_2$AR in whole cells**

**a, Data**

| Receptor | CGP12177 ($K_D$; nM) [*n*] | CGP20712A (log $K_D$) | *n* | NDD713 (log $K_D$) | *n* | NDD825 (log $K_D$) | *n* |
|---|---|---|---|---|---|---|---|
| hβ$_1$AR | 0.28 † | −8.97 ± 0.05 | 6 | −7.91 ± 0.03 | 7 | −8.32 ± 0.04 | 7 |
| hβ$_1$AR I118H | 0.21 # | −8.44 ± 0.06 | 6 | −7.29 ± 0.03 | 7 | −7.74 ± 0.04 | 7 |
| hβ$_1$AR V119I | 0.30 # | −9.11 ± 0.02 | 6 | −7.89 ± 0.04 | 7 | −7.98 ± 0.04 | 7 |
| hβ$_1$AR V189T | 0.38 ± 0.02 [*9*] | −8.99 ± 0.02 | 6 | −7.95 ± 0.03 | 7 | −8.30 ± 0.03 | 7 |
| hβ$_1$AR F359Y | 0.51 † | −8.63 ± 0.03 | 6 | −7.56 ± 0.03 | 7 | −7.95 ± 0.03 | 7 |
| hβ$_1$AR V360I | 0.34 ± 0.02 [*6*] | −8.88 ± 0.09 | 6 | −8.07 ± 0.05 | 7 | −8.34 ± 0.04 | 7 |
| hβ$_1$AR•hβ$_2$OBS | 1.048 ± 0.03 [*6*] | −7.78 ± 0.04 | 6 | −6.56 ± 0.03 | 7 | −6.92 ± 0.03 | 7 |
| hβ$_2$AR | 0.18 † | −6.01 ± 0.02 | 6 | −5.38 ± 0.02 | 7 | −5.89 ± 0.04 | 7 |
| hβ$_2$AR H93I | 0.30 # | −6.77 ± 0.02 | 6 | −6.39 ± 0.04 | 7 | −6.55 ± 0.03 | 7 |
| hβ$_2$AR I94V | 0.21 # | −6.16 ± 0.03 | 6 | −5.54 ± 0.02 | 7 | −6.19 ± 0.03 | 7 |
| hβ$_2$AR T164V | 0.21 ± 0.02 [*7*] | −5.80 ± 0.03 | 6 | −4.98 ± 0.04 | 7 | −5.48 ± 0.03 | 7 |
| hβ$_2$AR Y308F | 0.16 † | −6.02 ± 0.03 | 6 | −5.51 ± 0.02 | 7 | −5.94 ± 0.02 | 7 |
| hβ$_2$AR I309V | 0.23 † | −5.91 ± 0.02 | 6 | −5.46 ± 0.04 | 7 | −5.82 ± 0.04 | 7 |
| hβ$_2$AR•hβ$_1$OBS | 0.12 ± 0.01 [*6*] | −6.67 ± 0.03 | 6 | −6.47 ± 0.02 | 7 | −6.71 ± 0.02 | 7 |

Values are log $K_D$ (M), mean ± SEM of *n* independent experiments. Mutants are named by the parent receptor residue, its position in the human sequence, and the mutated residue; hβ$_1$AR•hβ$_2$OBS and hβ$_2$AR•hβ$_1$OBS carry all five substitutions. † values from Baker *et al.*[22]. # values from Lim *et al.*[23]

**b, *P* values**

| Receptor | CGP20712A | NDD713 | NDD825 |
|---|---|---|---|
| hβ$_1$AR | – | – | – |
| hβ$_1$AR I118H$^{2.64}$ | $2 \times 10^{-7}$ | $1 \times 10^{-12}$ | $2 \times 10^{-12}$ |
| hβ$_1$AR V119I$^{2.65}$ | 0.4 | 1.0 | $9 \times 10^{-7}$ |
| hβ$_1$AR V189T$^{4.56}$ | 1.0 | 1.0 | 1.0 |
| hβ$_1$AR F359Y$^{7.35}$ | $5 \times 10^{-4}$ | $2 \times 10^{-7}$ | $1 \times 10^{-7}$ |
| hβ$_1$AR V360I$^{7.36}$ | 0.9 | 0.04 | 1.0 |
| hβ$_1$AR•hβ$_2$OBS | $< 1 \times 10^{-15}$ | $1 \times 10^{-12}$ | $1 \times 10^{-12}$ |
| hβ$_2$AR | – | – | – |
| hβ$_2$AR H93I$^{2.64}$ | $< 1 \times 10^{-15}$ | $1 \times 10^{-12}$ | $1 \times 10^{-12}$ |
| hβ$_2$AR I94V$^{2.65}$ | $5 \times 10^{-3}$ | $9 \times 10^{-3}$ | $5 \times 10^{-7}$ |
| hβ$_2$AR T164V$^{4.56}$ | $4 \times 10^{-5}$ | $2 \times 10^{-10}$ | $2 \times 10^{-10}$ |
| hβ$_2$AR Y308F$^{7.35}$ | 1.0 | 0.06 | 0.9 |
| hβ$_2$AR I309V$^{7.36}$ | 0.1 | 0.5 | 0.7 |
| hβ$_2$AR•hβ$_1$OBS | $< 1 \times 10^{-15}$ | $1 \times 10^{-12}$ | $1 \times 10^{-12}$ |

*P* values from one-way ANOVA with post hoc Tukey analysis (see methods) with *P* values quoted for comparison of each ligand for parent receptor with each of the mutants (i.e. hβ1AR with hβ1-I118H, hβ1AR with hβ1-V119I etc from within the single analysis. *P* values ≤ $1 \times 10^{-3}$ are considered significant.

**Extended Data Table 4 | Equilibrium and kinetic binding parameters of CGP20712A, NDD713 and NDD825 at wild-type and chimeric β-adrenoceptors**

#### a, Data

| Ligand | Parameter | h$β_1$AR | h$β_2$AR | h$β_1AR_{out}$/h$β_2AR_{in}$ | h$β_2AR_{out}$/h$β_1AR_{in}$ |
|---|---|---|---|---|---|
| CGP20712A | log $K_D$ (M) | −8.19 ± 0.02 (6) | −4.95 ± 0.04 (5) | −6.23 ± 0.02 (5) | −5.29 ± 0.01 (4) |
| | log $k_{on}$ ($M^{-1}$ $min^{-1}$) | 7.17 ± 0.04 (6) | 5.34 ± 0.09 (5) | 6.26 ± 0.06 (5) | 5.38 ± 0.05 (4) |
| | log $k_{off}$ ($min^{-1}$) | −1.02 ± 0.04 (6) | 0.39 ± 0.08 (5) | 0.03 ± 0.05 (5) | 0.09 ± 0.05 (4) |
| NDD713 | log $K_D$ (M) | −7.93 ± 0.03 (6) | −5.02 ± 0.03 (6) | −5.96 ± 0.02 (6) | −5.03 ± 0.01 (6) |
| | log $k_{on}$ ($M^{-1}$ $min^{-1}$) | 7.39 ± 0.03 (6) | 4.86 ± 0.03 (6) | 5.77 ± 0.04 (6) | 4.82 ± 0.04 (6) |
| | log $k_{off}$ ($min^{-1}$) | −0.54 ± 0.01 (6) | −0.16 ± 0.03 (6) | −0.19 ± 0.05 (6) | −0.20 ± 0.05 (6) |
| NDD825 | log $K_D$ (M) | −8.00 ± 0.02 (6) | −5.00 ± 0.01 (3) | −5.87 ± 0.02 (4) | −5.46 ± 0.01 (4) |
| | log $k_{on}$ ($M^{-1}$ $min^{-1}$) | 7.14 ± 0.01 (6) | 5.79 ± 0.10 (3) | 6.40 ± 0.09 (5) | 5.90 ± 0.14 (6) |
| | log $k_{off}$ ($min^{-1}$) | −0.86 ± 0.01 (6) | 0.79 ± 0.09 (3) | 0.44 ± 0.03 (4) | 0.23 ± 0.04 (4) |

Values are mean ± SEM of log-transformed replicate values; *n* (replicates) in parentheses; all values are from three independent experiments each performed in duplicate. $K_D$, equilibrium dissociation constant; $k_{on}$, association rate constant; $k_{off}$, dissociation rate constant. $K_D$ was calculated for each replicate as $k_{off}/k_{on}$ and then averaged, so log $K_D$ is not necessarily the difference between the tabulated mean log $k_{off}$ and log $k_{on}$ where the three parameters derive from different numbers of replicates.

#### b, *P* values

| Ligand | Parameter | h$β_1$AR vs h$β_2$AR | h$β_1$AR vs h$β_1AR_{out}$/h$β_2AR_{in}$ | h$β_1$AR vs h$β_2AR_{out}$/h$β_1AR_{in}$ | h$β_2$AR vs h$β_1AR_{out}$/h$β_2AR_{in}$ | h$β_2$AR vs h$β_2AR_{out}$/h$β_1AR_{in}$ |
|---|---|---|---|---|---|---|
| CGP20712A | log $K_D$ | $1.9 \times 10^{-14}$ | $6.1 \times 10^{-14}$ | $2.5 \times 10^{-14}$ | $1.1 \times 10^{-9}$ | $7.9 \times 10^{-5}$ |
| | log $k_{on}$ | $1.0 \times 10^{-8}$ | $4.6 \times 10^{-7}$ | $3.3 \times 10^{-9}$ | $2.2 \times 10^{-5}$ | 0.71 |
| | log $k_{off}$ | $3.0 \times 10^{-8}$ | $4.6 \times 10^{-8}$ | $9.8 \times 10^{-8}$ | 0.0051 | 0.019 |
| NDD713 | log $K_D$ | $3.3 \times 10^{-15}$ | $3.0 \times 10^{-14}$ | $2.9 \times 10^{-16}$ | $3.3 \times 10^{-11}$ | 0.756 |
| | log $k_{on}$ | $9.4 \times 10^{-14}$ | $4.6 \times 10^{-11}$ | $6.8 \times 10^{-13}$ | $7.0 \times 10^{-9}$ | 0.501 |
| | log $k_{off}$ | $2.9 \times 10^{-7}$ | $2.0 \times 10^{-5}$ | $3.6 \times 10^{-5}$ | 0.606 | 0.416 |
| NDD825 | *log* $K_D$ | $6.4 \times 10^{-13}$ | $3.4 \times 10^{-13}$ | $4.2 \times 10^{-14}$ | $1.5 \times 10^{-7}$ | $6.6 \times 10^{-7}$ |
| | log $k_{on}$ | $1.6 \times 10^{-7}$ | $1.3 \times 10^{-5}$ | $4.5 \times 10^{-6}$ | 0.0058 | 0.613 |
| | log $k_{off}$ | $2.8 \times 10^{-8}$ | $8.5 \times 10^{-11}$ | $1.2 \times 10^{-9}$ | 0.010 | 0.0017 |

*P* values from one-way ANOVA with post hoc Tukey analysis (see methods). *P* values ≤ 1 x $10^{-3}$ are considered significant.

**Extended Data Table 5 | Details of all atomistic molecular dynamics simulations run. Note that all simulations used a membrane composition of 80% POPC and 20% cholesterol**

| Simulation type | System | Source | Box size (nm) | Simulation length (ns) | Number of reps/windows/ walkers |
|---|---|---|---|---|---|
| Unbiased MD | Inactive h$\beta_1$AR | PDB 7BVQ[25] | 7.7 x 7.7 x 11.0 | 500 | 5 |
| | Inactive h$\beta_2$AR | PDB 2RH1[43] | 7.7 x 7.7 x 12.0 | 500 | 5 |
| Umbrella MD | Inactive h$\beta_1$AR | PDB 7BVQ[25] | 7.5 x 7.5 x 11.6 | 20 | 16 |
| | Inactive h$\beta_2$AR | PDB 2RH1[43] | 7.4 x 7.4 x 12.8 | 20 | 16 |
| mwWT-MetaD | Inactive h$\beta_1$AR + CGP20712A | PDB 7BVQ[25] | 7.1 x 7.1 x 13.5 | 926.8 | 16 |
| | Inactive h$\beta_2$AR + CGP20712A | PDB 2RH1[43] | 7.2 x 7.2 x 14.1 | 891.6 | 16 |
